# APEC: an accesson-based method for single-cell chromatin accessibility analysis

**DOI:** 10.1101/646331

**Authors:** Bin Li, Young Li, Kun Li, Lianbang Zhu, Qiaoni Yu, Pengfei Cai, Jingwen Fang, Wen Zhang, Pengcheng Du, Chen Jiang, Kun Qu

## Abstract

The development of sequencing technologies has promoted the survey of genome-wide chromatin accessibility at single-cell resolution; however, comprehensive analysis of single-cell epigenomic profiles remains a challenge. Here, we introduce an accessibility pattern-based epigenomic clustering (APEC) method, which classifies each individual cell by groups of accessible regions with synergistic signal patterns termed “accessons”. By integrating with other analytical tools, this python-based APEC package greatly improves the accuracy of unsupervised single-cell clustering for many different public data sets. APEC also predicts gene expressions, identifies significant differential enriched motifs, discovers super enhancers, and projects pseudotime trajectories. Furthermore, we adopted a fluorescent tagmentation-based single-cell ATAC-seq technique (ftATAC-seq) to investigated the per cell regulome dynamics of mouse thymocytes. Associated with ftATAC-seq, APEC revealed a detailed epigenomic heterogeneity of thymocytes, characterized the developmental trajectory and predicted the regulators that control the stages of maturation process. Overall, this work illustrates a powerful approach to study single-cell epigenomic heterogeneity and regulome dynamics.

## INTRODUCTION

As a technique for probing genome-wide chromatin accessibility in a small number of cells *in vivo*, the assay for transposase-accessible chromatin with high-throughput sequencing (ATAC-seq) has been widely applied to investigate the cellular regulomes of many important biological processes ^1^, such as hematopoietic stem cell (HSC) differentiation ^2^, embryonic development ^3^, neuronal activity and regeneration ^4, 5^, tumor cell metastasis^6^, and patient responses to anticancer drug treatment ^7^. Recently, several experimental schemes have been developed to capture chromatin accessibility at single-cell/nucleus resolution, i.e., single-cell ATAC-seq (scATAC-seq) ^8^, single-nucleus ATAC-seq (snATAC-seq) ^9, 10^, and single-cell combinatorial indexing ATAC-seq (sci-ATAC-seq) ^11, 12^, which significantly extended researchers’ ability to uncover cell-to-cell epigenetic variation and other fundamental mechanisms that generate heterogeneity from identical DNA sequences. By contrast, the in-depth analysis of single-cell chromatin accessibility profiles for this purpose remains a challenge. Numerous efficient algorithms have been developed to accurately normalize, cluster and visualize cells from single-cell transcriptome sequencing profiles, including but not limited to Seurat ^13^, SC3 ^14^, SIMLR ^15^, and SCANPY^16^. However, most of these algorithms are not directly compatible with a single-cell ATAC-seq dataset, for which the signal matrix is much sparser. To characterize scATAC-seq data, the Greenleaf lab developed an algorithm named chromVAR ^17^, which aggregates mapped reads at accessible sites based on annotated motifs of known transcription factors (TFs) and thus projects the sparse per accessible peak per cell matrix to a bias-corrected deviation per motif per cell matrix and significantly stabilizes the data matrix for downstream clustering analysis. Other mathematical tools, such as the latent semantic indexing (LSI) ^11, 12^, Cicero ^18^, cisTopic ^19^, and SnapATAC ^20^ have also been applied to process single-cell/nucleus ATAC-seq data ^10, 12^. However, great challenges still remain for current algorithms to precisely cluster large number of cells and predict gene expressions from single cell chromatin accessibility profiles. Therefore, a refined algorithm is needed to better categorize cell subgroups with minor differences, thereby providing a deeper mechanistic understanding of single-cell epigenetic heterogeneity and regulation.

Here, we introduce a new single-cell chromatin accessibility analysis toolkit named APEC (accessibility pattern-based epigenomic clustering), which combines peaks with the same signal fluctuation among all single cells into peak groups, termed “accessons”, and converts the original sparse cell-peak matrix to a much denser cell-accesson matrix for cell type categorization (Fig. 1a). In contrast to previous methods, this accesson-based reduction scheme naturally groups synergistic accessible regions genome-wide together without a priori knowledge of genetic information (such as TF motifs or genomic distance) and provides an efficient, accurate, robust and rapid cell clustering from single-cell ATAC-seq profiles. APEC was also integrated into a head-to-toe program package that has been made available on GitHub (https://github.com/QuKunLab/APEC).

**Figure 1.**
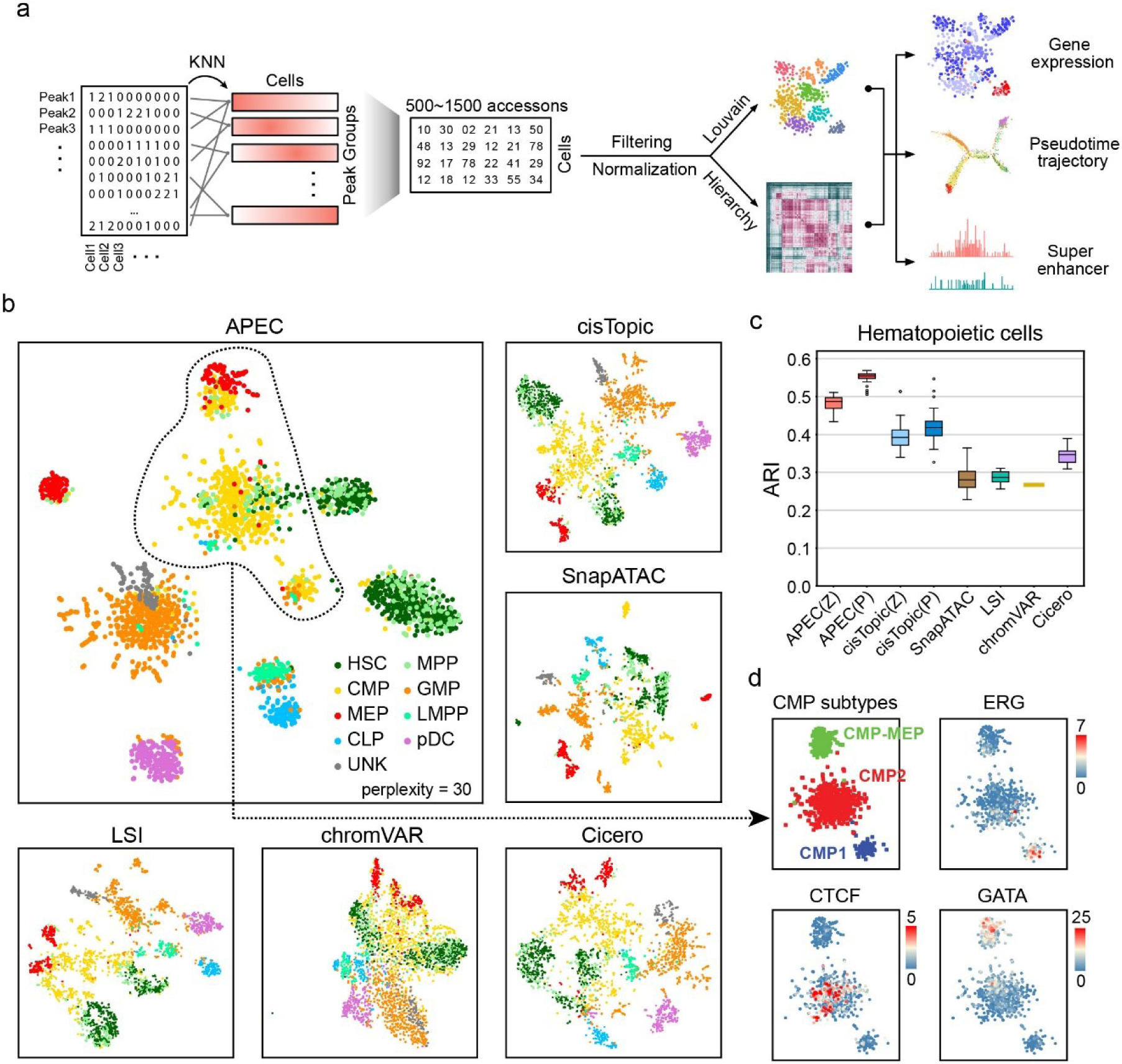
The accesson matrix constructed from the sparse fragment count matrix improved the clustering of scATAC-seq data. (**a**) Step-by-step workflow of APEC. Peaks were grouped into accessons by their accessibility pattern among cells with the K nearest neighbors (KNN) method. (**b**) t-Distributed Stochastic Neighbor Embedding (tSNE) diagrams of the hematopoietic single cells dataset based on the dimension-transformed matrices from different algorithms, i.e., APEC: accesson matrix; cisTopic: topic matrix; LSI: LSI matrix; chromVAR: bias-corrected deviation matrix; Cicero: aggregated model matrix. The cells are FACS-indexed human hematopoietic cells, including HSCs (hematopoietic stem cells), MPPs (multipotent progenitors), LMPPs (lymphoid-primed multipotential progenitors), CMPs (common myeloid progenitors), CLPs (common lymphoid progenitors), pDCs (plasmacytoid dendritic cells), GMPs (granulocyte-macrophage progenitors), MEPs (megakaryocyte-erythroid progenitors), and UNK (unknown type) cells. (**c**) The ARI (Adjusted Rand Index) values for the clustering of the human hematopoietic cells by different algorithms. Same as the two normalization methods applied in cisTopic, we normalized the accesson matrix in APEC based on probability (P) and z-score (Z). Center line, median; box limits, upper and lower quartiles; whiskers, 1.5x interquartile range; points, outliers. (**d**) Three CMP subtypes identified in APEC and the motifs enriched in each cell subtype.

## RESULTS

### Accesson-based algorithm improves single-cell clustering

To test the performance of APEC, we first obtained data from previous publications that performed scATAC-seq on a variety of cell types with known identity during hematopoietic stem cell (HSC) differentiation ^21^ as a gold standard. Compared to other state-of-the-art single cell chromatin accessibility analysis methods, this new accesson-based algorithm can clearly cluster cells into their corresponding identities according to the Adjusted Rand Index (ARI) (Fig. 1b & c). Here, we adopted the Louvain clustering algorithm for all the methods, and assumed that the number of cell types was unknown to ensure that the clustering analysis was unsupervised. To calculate ARI values, we ignored all “unknown” cells, and sampled the tunable parameter (e.g., accesson number in APEC, random seed in cisTopic, etc.) multiple times for each tool (see Methods). On average, 67% of cells were correctly classified by APEC with ARI=0.483/0.552 (using matrices normalized by Z-score or probability, respectively), while cisTopic was the second most accurate method to predict cell identities (average ARI=0.392/0.418) (Supplementary Table 1).

Moreover, APEC identified 3 sub-clusters of CMP cells that were not discovered by any other algorithms, namely CMP1, CMP2 and CMP-MEP (Fig. 1d). CMP1 cells are early stage of CMPs that enriched TFs associated with stem cell self-renewal, such as Erg ^22^; CMP2 cells are enriched with CTCF motif, suggesting that these cells are at the fate decision stage with CTCF associated chromatin remodeling ^23^; CMP-MEP cells are considered as MEP committed CMPs, and are strongly enriched with crucial regulators for MEP differentiation, such as GATA1 ^24^. However, these 3 CMP sub-clusters were not as clearly distinguishable and cells were more scattered on the tSNE maps generated by other methods (Supplementary Fig. 1). More details about the distribution of these 3 sub-clusters of CMP cells on the development trajectory will be discussed later in the section of pseudotime prediction.

To further confirm the superiority of APEC, we performed the same comparison analysis with another scATAC-seq dataset on three distinct cell types, namely, lymphoid-primed multipotent progenitors (LMPPs), monocytes, and HL-60 lymphoblastoid cells (HL60), and four similar cell types, namely, blast cells and leukemic stem cells (LSCs) from two acute myeloid leukemia (AML) patients ^17^. We found that both APEC and cisTopic were tied for best to classify these cells (Supplementary Fig. 2a). Interestingly, APEC, cisTopic and LSI were all capable of almost perfectly separating the three distinct cell types (LMPPs, monocytes, and HL60), with ARI = 1.000/0.988, 0.987/0.987 and 0.969, respectively. However, in terms of clustering the four similar cell types from AML patients, APEC (average ARI=0.575/0.564) outperformed other tools (Supplementary Fig. 2b), suggesting that APEC was the most sensitive among all the tools. Since each method can generate varying numbers of clusters depending on the parameters used, we benchmarked the performance of all the methods using ARI across a wide range of tunable parameters to ensure the reliability of their predictions (Supplementary Fig. 2c, Supplementary Table 2). To further test the robustness of APEC at low sequencing depth, we randomly selected reads from the raw data and calculated the ARI values for each down-sampled data. APEC exhibits better performance at sequencing depths as low as 20% of the original data (Supplementary Fig. 2d), confirming the sensitivity of the algorithm.

Compare with chromVAR, the contribution of the minor differences between similar cells is aggregated in accessons but diluted in motifs. For example, APEC identified prominent super-enhancers around the genes *N4BP1* ^25^ and *GPHN* ^26^ in the LSC cells from AML patient 1 (P1-LSC) but not the other cell types (Supplementary Fig. 3a & b). These two loci were also confirmed as super enhancers by ROSE ^27^ (the top 2 candidates in Supplementary Table 3). We noticed that all peaks in these super-enhancers were classified into one accesson that was critical for distinguishing P1-LSCs from P2-LSCs, P1-blast cells and P2-blast cells. However, these peaks were distributed in multiple TF motifs, which significantly diluted the contributions of the minor differences (Supplementary Fig. 3c & d).

In contrast to Cicero, which aggregates peaks based on their cis-co-accessibilities networks (CCAN) within a certain range of genomic distance ^18^, APEC combine synergistic peaks genome-wide. Take the human hematopoietic cells dataset as an example, 600 accessons were built from the 54,212 peaks, and each accesson contained ∼40 peaks (median number) compare with ∼4 peaks in each CCAN (Supplementary Fig. 4a & b). The average distance between peaks in a same accesson is ∼50 million base pairs (compared with ∼0.2 million bps from CCAN), and over 57% of accessons contain peaks from more than 15 different chromosomes. From the same dataset, Cicero identified 732,306 pairs of site links from 25,102 peaks, and information from the remaining peaks were simply discarded. APEC identified more than 9.2 million pairs of site links from all the 54,212 peaks, within which only 3080 site links were identified by both methods (Supplementary Fig. 4c), therefore, APEC and Cicero are two completely different approaches. Furthermore, Buenrostro et al. showed that the covariation of the accessible sites across all the cells may reflect the spatial distance between the corresponding peaks ^8^. By integrating the chromatin conformation profiles from Hi-C experiments with the scATAC-seq profile for the same cells, we found that peaks in the same accesson are spatially much closer to each other than randomly selected peaks (Supplementary Fig. 5a, P-value<10^-7^), suggesting that they may belong to the same topologically associated domains (TADs) (Supplementary Fig. 5b).

Speed and scalability are now extremely important for single-cell analytical tools due to the rapid growth in the number of cells sequenced in each experiment. We benchmarked APEC and all the other tools based on a random sampling of the mouse *in vivo* single-cell chromatin accessibility atlas dataset ^28^, which contains 81,173 high quality cells. Taking into account of all the 436,206 peaks, it took APEC 310 min to cluster 80,000 cells with 1 CPU thread (Supplementary Fig. 5c). We also randomly select 100,000 peaks from the entire dataset to test the computer time spent by these tools (Supplementary Fig. 5d). In addition, APEC is very stable for a wide range of parameter values used in the algorithm, such as the number of accessons, nearest neighbors and principle components (Supplementary Fig. 5e-g).

### APEC is applicable to other single-cell chromatin detection techniques

To evaluate the compatibility and performance of APEC with other single-cell chromatin accessibility detection techniques, such as snATAC-seq ^10^, transcript-indexed scATAC-seq ^29^ and sciATAC-seq ^11^, APEC was also tested with the data sets generated by those experiments. For example, APEC discovered 14 cell subpopulations in adult mouse forebrain snATAC-seq data ^10^, including four clusters of excitatory neurons (EX1-5), five groups of inhibitory neurons (IN1-5), astroglia cells (AC1&2), oligodendrocyte cells (OC), and microglial cells (MG; Fig. 2a & b). To quantify gene expression level, we defined a gene’s score as the average signal of the peaks close to its TSS region (Fig. 2c; see Methods). With that, we identified 5 excitatory subpopulations and 5 distinct inhibitory subpopulations, and all the cell groups were clearly distinguished from each other by hierarchical clustering (Fig. 2d). In contrast, more than 29.7% (946 out of 3034) of cells were unable to be correctly assigned into any subpopulation of interest in previous study^10^. Besides, scores for several marker genes such as *Neurod6* and *Aldh1l1* were not available in Cicero and cisTopic, respectively, making it difficult to identify the corresponding cell clusters (i.e. c0∼c2 in Supplementary Fig. 6a & b). The SnapATAC algorithms however, mis-clustered the AC1/2 with excitatory neurons in the correlation matrix (Supplementary Fig. 6c). On the other hand, the motif enrichment analysis module in APEC identified cell type-specific regulators that are also consistent with previous publications ^10^. For example, the NEUROD1 and OLIG2 motifs were generally enriched on excitatory clusters (EX1\2\3\5); the MEF2C motif was more enriched on EX1/3/4/5; the motifs of MEIS2 and DLX2 were differentially enriched on different subtypes of inhibitory neurons (IN3 and IN2/4, respectively); and the NOTO, SOX2, and ETS1 motifs were enriched on the AC1, OC, and MG clusters, respectively (Fig. 2e). These results confirm that APEC can distinguish and categorize single cells with great sensitivity and reliability.

**Figure 2.**
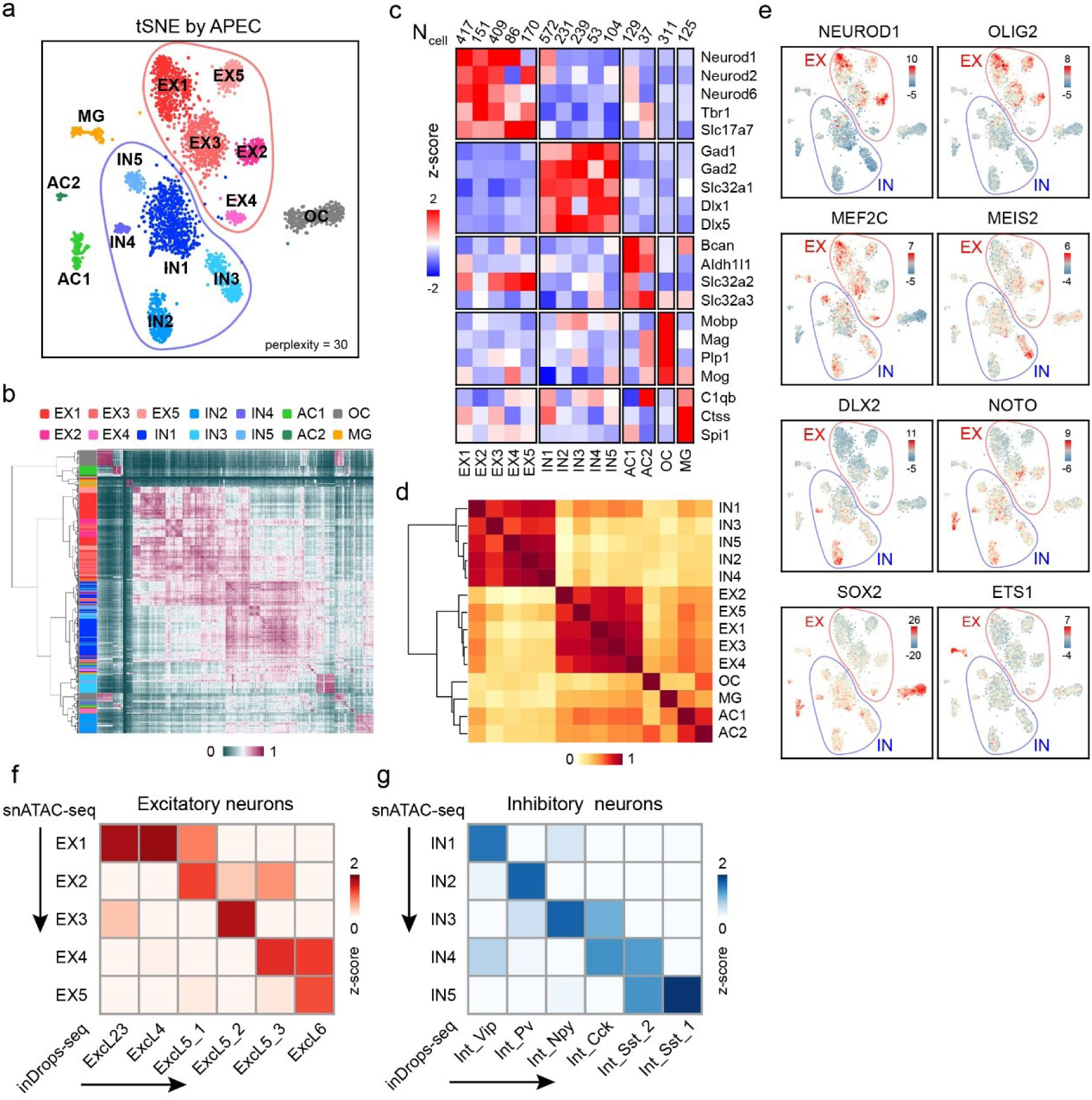
APEC improved the cell type classification of adult mouse forebrain snATAC-seq data. (**a**) A tSNE diagram demonstrates the APEC clustering of forebrain cells. (**b**) Hierarchical clustering of the cell-cell correlation matrix. The side bar denotes cell clusters from APEC. (**c**) Average scores of the marker genes for each cell cluster generated by the method mentioned in the data source paper ^10^, and normalized by the standard score (z-score). The top row lists the number of cells in each cluster. (**d**) Hierarchical clustering of the cluster-cluster correlation matrix. (**e**) Differential enrichments of cell type-specific motifs in each cluster. (**f, g**) Fisher’s exact test between the differential genes of the excitatory (EX) and inhibitory (IN) neurons from snATAC-seq and the signature genes of the excitatory (Excl) and inhibitory (Int) neurons defined in inDrops-seq^30^. The heatmaps show the –log(P-value) normalized by calculating the z-scores through each row and column.

Since single-cell transcriptome analysis is also capable to identify novel cell subpopulations, it is critical to anchor the cell types identified from scATAC-seq to those from scRNA-seq. Hrvatin et al. identified multiple excitatory and inhibitory neuronal subtypes in the mouse visual cortex using single cell inDrops sequencing^30^ and provided top 20 signature genes that distinguished these cell subtypes. However, due to the sparseness of snATAC-seq matrix, scores of many signature genes were not strong enough to distinguish the cell sub-clusters. To overcome this, we developed a gene sets overlap algorithm to associate cell clusters from scATAC-seq and scRNA-seq profiles (see Methods). We found that sub-cluster EX1∼5 and IN1∼5 in snATAC-seq can nicely correspond to the neuron subtypes classified by Hrvatin et al (Fig. 2f-g). These results highlight the potential advantages of the accesson-based approach for the integrative analysis of scATAC-seq and scRNA-seq data.

### APEC constructs a pseudotime trajectory that predicts cell differentiation lineage

Cells are not static but dynamic entities, and they have a history, particularly a developmental history. Although single-cell experiments often profile a momentary snapshot, a number of remarkable computational algorithms have been developed to pseudo-order cells based on the different points they were assumed to occupy in a trajectory, thereby leveraging biological asynchrony ^31, 32^. For instance, Monocle ^32, 33^ constructs the minimum spanning tree, and Wishbone ^34^ and SPRING ^35^ construct the nearest neighbor graph from single-cell transcriptome profiles. These tools have been widely used to depict neurogenesis ^36^, hematopoiesis ^37, 38^ and reprogramming ^39^. APEC integrates the Monocle algorithm into the accesson-based method and enables pseudotime prediction from scATAC-seq data ^21^ and was applied to investigate HSC differentiation linages (Fig. 3a). Principal component analysis (PCA) of the accesson matrix revealed multiple stages of the lineage during HSC differentiation (Fig. 3b) and was consistent with previous publications ^2, 21^. After utilizing the Monocle package, APEC provided more precise pathways from HSCs to the differentiated cell types (Fig. 3c). In addition to the differentiation pathways to MEP cells through the CMP state and to CLP cells through the LMPP state, MPP cells may differentiate into GMP cells through two distinct trajectories: Path A through the CMP state and Path B through the LMPP state, which is consistent with the composite model of HSC and blood lineage commitment^40^. Notably, pDCs from bone marrow are CD34^+^ (Supplementary Fig. 7), indicative of precursors of pDCs. APEC suggested that pDC precursors were derived from CLP cells on the pseudotime trajectory (Fig. 3c), which also agrees with previous reports ^41^. Furthermore, APEC incorporated the chromVAR algorithm to determine the regulatory mechanisms during HSC differentiation by evaluating the deviation of each TF along the single-cell trajectory. As expected, the HOX motif is highly enriched in the accessible sites of HSCs/MPP cells, as are the GATA1, CEBPB and TCF4 motifs, which exhibit gradients that increase along the erythroid, myeloid and lymphoid differentiation pathways, respectively ^21^ (Fig. 3d). We also noticed that the TF regulatory strategies of the two paths from MPP towards GMP cells were very different. In addition, the 3 CMP sub-clusters identified in Figure 1 were differentially distributed along the developmental trajectory (Fig. 3e). CMP1 cells that close to HSCs and MPPs are early stage CMPs; CMP2 cells are distributed in both the GMP and MEP branches; CMP-MEP cells are MEP committed CMPs and are dominantly distributed in the MEP differentiation branch. The distributions of these CMP sub-clusters are also consistent with the functions of their enriched motifs mentioned in the first section (Fig. 1d) ^22–24^. Finally, we generated a hematopoiesis tree based on the APEC analysis (Fig. 3f).

**Figure 3.**
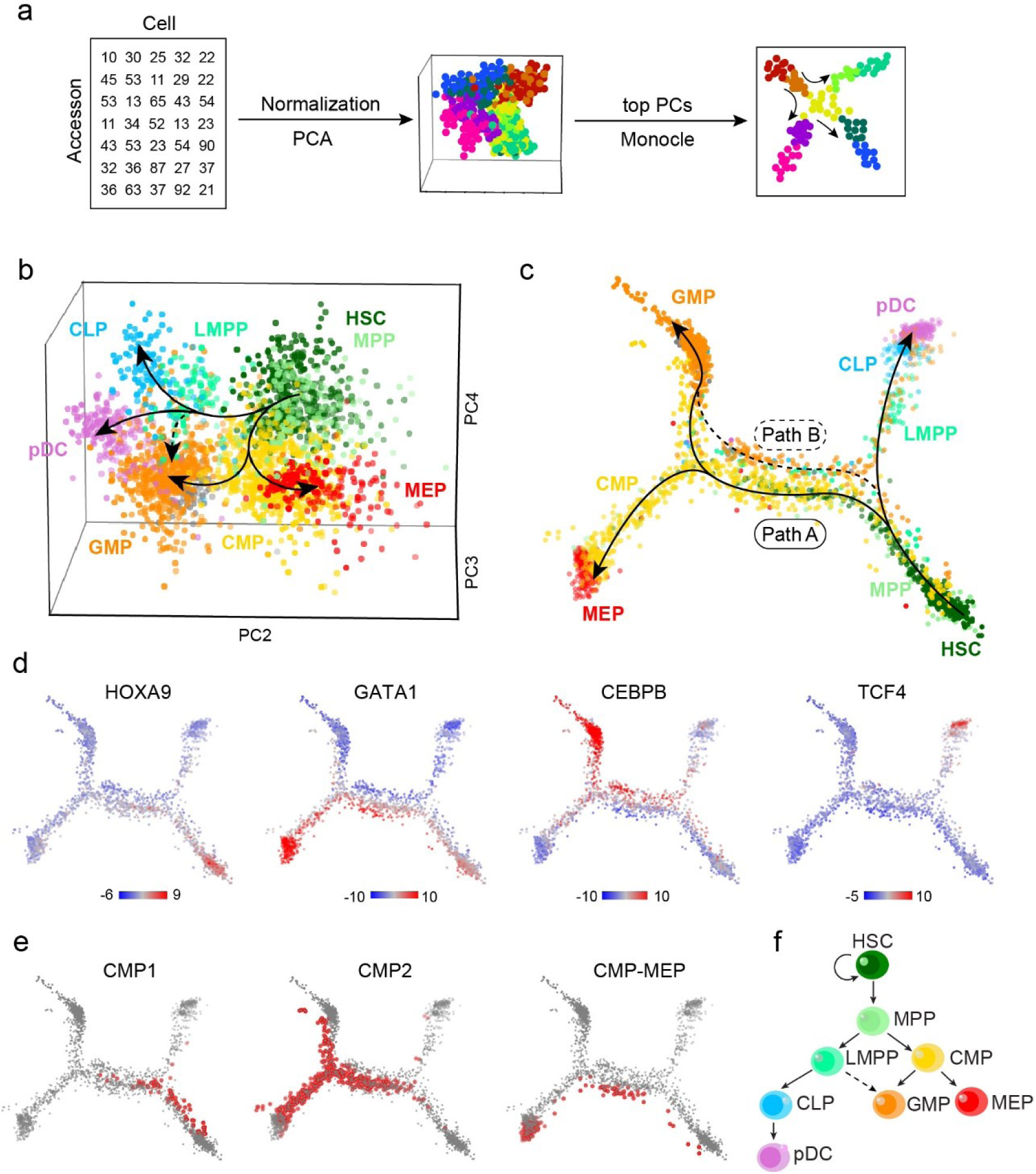
APEC constructed a differentiation pathway from scATAC-seq data from human hematopoietic cells. (**a**) The pseudotime trajectory construction scheme based on the accesson matrix and Monocle. (**b**) Principal component analysis (PCA) of the accesson matrix for human hematopoietic cells. The first principal component is not shown here because it was highly correlated with sequencing depth ^21^. HSC, hematopoietic stem cell; MPP, multipotent progenitor; LMPP, lymphoid-primed multipotential progenitor; CMP, common myeloid progenitor; CLP, common lymphoid progenitor; pDC, plasmacytoid dendritic cell; GMP, granulocyte-macrophage progenitor; MEP, megakaryocyte-erythroid progenitor. (**c**) Pseudotime trajectory for the same data constructed by applying Monocle on the accesson matrix. Paths A and B represent different pathways for GMP cell differentiation. (**d**) The deviations of significant differential motifs (HOXA9, GATA1, CEBPB, and TCF4) plotted on the pseudotime trajectory. (**e**) Distributions of the CMP sub-clusters on the trajectory. (**f**) Modified schematic of human hematopoietic differentiation.

Furthermore, we benchmarked the performance of APEC and of all the other tools in constructing a pseudotime trajectory from the scATAC-seq profile on the same dataset. We found that (1) when the raw peak count matrix was invoked into Monocle, almost none developmental pathways were constructed (Supplementary Fig. 8a), suggesting that the peak aggregation step in APEC greatly improves the pseudotime estimation; (2) APEC + Monocle provides the most precise pathways from HSCs to differentiated cells, compared to other methods, including cisTopic, SnapATAC, LSI, Cicero, and chromVAR (Supplementary Fig. 8b-f); and (3) when we applied other pseudotime trajectory construction methods, such as SPRING ^35^, after APEC, a similar though less clear cell differentiation diagram was also obtained, suggesting the reliability of our prediction (Supplementary Fig. 8g).

### APEC reveals the single-cell regulatory heterogeneity of thymocytes

T cells generated in the thymus play a critical role in the adaptive immune system, and the development of thymocytes can be divided into 3 main stages based on the expression of the surface markers CD4 and CD8, namely, CD4 CD8 double-negative (DN), CD4 CD8 double-positive (DP) and CD4 or CD8 single-positive (CD4SP or CD8SP, respectively) stages^42^. However, due to technical limitations, our genome-wide understanding of thymocyte development at single-cell resolution remains unclear. Typically, more than 80% of thymocytes stay in the DP stage in the thymus, whereas DN cells account for only approximately 3% of the thymocyte population. To eliminate the impacts of great differences in proportion, we developed a fluorescent tagmentation- and FACS-sorting-based scATAC-seq strategy (ftATAC-seq), which combined the advantages of ATAC-see ^43^ and Pi-ATAC-seq ^44^ to manipulate the desired number of target cells by indexed sorting (Fig. 4a). Tn5 transposomes were fluorescently labeled in each cell to evaluate the tagmentation efficiency so that cells with low ATAC signals could be gated out easily (Fig. 4b, Supplementary Fig. 9a). With ftATAC-seq, we acquired high-quality chromatin accessibility data for 352 index-sorted DN, DP, CD4SP, and CD8SP single cells and 352 mixed thymocytes (Supplementary Fig. 9b-d). Correlation analysis with the published bulk ATACs-eq data of thymocytes ^45^ indicates that the cells we sorted in ftATAC-seq were correctly labeled (Supplementary Fig. 9e). We then applied APEC on this dataset to investigate the chromatin accessibility divergence during developmental process and to reveal refined regulome heterogeneity of mouse thymocytes at single-cell resolution. Taking into account of all the 130685 peaks called from the raw sequencing data, APEC aggregated 600 accessons and successfully assigned over 82% of index-sorted DN, DP, CD4SP and CD8SP cells into the correct subpopulations (Fig. 4c & 4d). As expected, the majority of randomly sorted and mixed thymocytes were classified into DP subtypes based on hierarchical clustering of cell-cell correlation matrix, which was consistent with the cellular subtype proportions in the thymus. APEC further classified all thymocytes into 11 subpopulations, including 2 DN, 6 DP, 1 CD4SP, 2 CD8SP, suggesting that extensive epigenetic heterogeneity exists among cells with the same CD4 and CD8 surface markers (Fig. 4e). For instance, there are four main subtypes of DN cells, according to the expression of the surface markers CD44 and CD25^46^, while two clusters were identified in ftATAC-seq. The accessibility signals around the *Il2ra* (Cd25) and *Cd44* gene loci demonstrated that DN.A1 comprised CD44^+^CD25^-^ and CD44^+^CD25^+^ DN subtypes (DN1 and DN2), and DN.A2 cells comprised CD44^-^CD25^+^ and CD44^-^CD25^-^ subtypes (DN3 and DN4), suggesting significant chromatin changes between DN2 and DN3 cell development (Fig. 4f).

**Figure 4.**
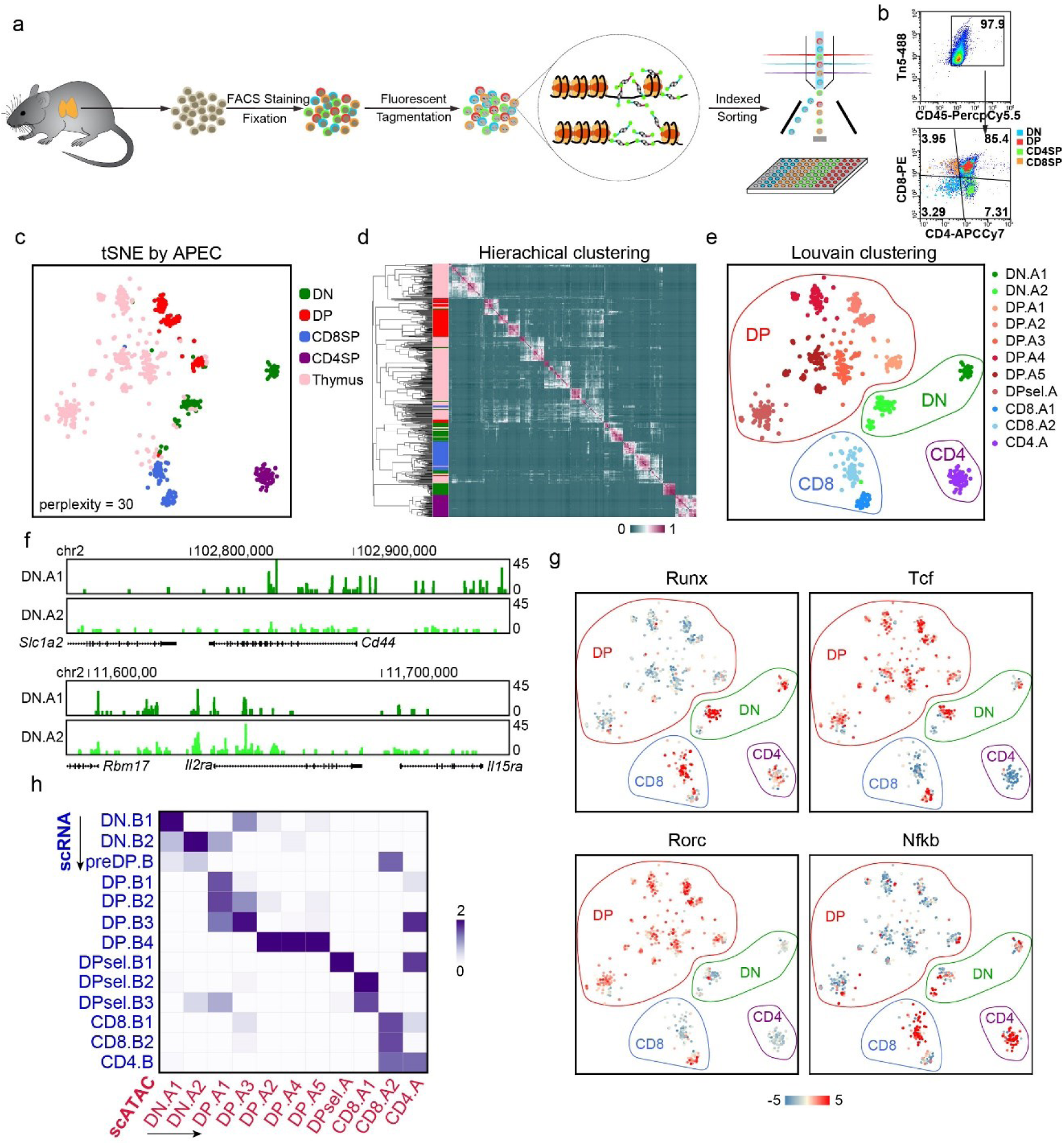
APEC accurately identified cell subtypes based on scATAC-seq data from *Mus musculus* thymocytes. (**a**) Experimental workflow of the fluorescent tagmentation- and FACS-sorting-based scATAC-seq strategy (ftATAC-seq). (**b**) Indexed sorting of double-negative (DN), double-positive (DP), CD4^+^ single-positive (CD4SP), and CD8^+^ single-positive (CD8SP) cells with strong tagmentation signals. (**c**) The tSNE of thymocyte single-cell ftATAC-seq data based on the accesson matrix, in which the cells are labeled by the sorting index. (**d**) Hierarchical clustering of the cell-cell correlation matrix. On the sidebar, each cell was colored by the sorting index. (**e**) The accesson-based Louvain method clustered thymocytes into 11 subtypes. DN.A1 (dark green) & A2 (light green), double-negative clusters; DP.A1∼A5 and DPsel.A, double-positive clusters; CD8.A1 (dark blue) & A2 (light blue), CD8^+^ single-positive clusters; CD4.A (purple), CD4^+^ single-positive cluster. (**f**) Average fragment counts of two DN clusters around the marker genes *Cd44* and *Il2ra*. (**g**) Differential enrichment of the motifs Runx, Tcf, Rorc, and Nfkb in the cell clusters. (**h**) Z-score of the Fisher exact test -log(p-value) of the common differential genes between the cell clusters from different experiments. The row and column clusters were identified by data from single-cell transcriptome (SMART-seq) and chromatin accessibility (ftATAC-seq) analysis respectively.

Many TFs have been reported to be essential in regulating thymocyte development, and we found that their motifs were remarkably enriched at different stages during the process (Fig. 4g). For instance, Runx3 is well known for regulating CD8SP cells ^47^, and we observed significant enrichment of the RUNX motif on DN cells and a group of CD8SP cells. Similarly, the TCF ^48, 49^, RORC ^50^ and NFkB ^51^ family in regulating the corresponding stages during this process. More enriched TF motifs in each cell subpopulation were also observed, suggesting significant regulatory divergence in thymocytes (Supplementary Fig. 10). Interestingly, two clusters of CD8SP cells appear to be differentially regulated based on motif analysis, in which CD8.A1 cells are closer to DP cells, while CD8.A2 cells are more distant at the chromatin level, suggesting that CD8.A2 cells are more mature CD8SP cells, and CD8.A1 cells are in a transitional state between DP and SP cells.

APEC is capable of integrating single-cell transcriptional and epigenetic information by scoring gene sets of interest based on their nearby peaks from scATAC-seq, thereby converting the chromatin accessibility signals to values that are comparable to gene expression profiles (see Methods). To test the performance of this integrative analysis approach and to evaluate the accuracy of thymocyte classification by APEC, we assayed the transcriptomes of single thymocytes and obtained 357 high-quality scRNA-seq profiles using the SMART-seq2 protocol ^52^. Unsupervised analysis of gene expression profiles clustered these thymocytes into 13 groups in Seurat ^13^ (Supplementary Fig. 11a & b), and each subpopulation was identified based on known feature genes (Supplementary Fig. 11c & d). We then adopted fisher’s exact test on the shared differential genes in cell clusters identified from scATAC-seq and scRNA-seq profiles (see Methods), and observed a strong correlation between the subtypes identified from the transcriptome and those from chromatin accessibility (Fig. 4h), confirming the reliability and stability of cellular classification using APEC.

### APEC reconstructs the thymocyte developmental trajectory

APEC is capable of constructing a pseudotime trajectory and then predicting the cell differentiation lineage from a “snapshot” of single-cell epigenomes (Fig. 3). We applied APEC to recapitulate the developmental trajectory and thereby reveal the single-cell regulatory dynamics during the maturation of thymocytes. Pseudotime analysis based on single-cell ATAC-seq data shaped thymocytes into 5 developing stages (Fig. 5a, Supplementary Fig. 12a & b), where most of the cells in stages 1, 2, 4, and 5 were DN, DP, CD8SP and CD4SP cells, respectively. APEC also identified a transitional stage 3, which was mainly consisted of last stages of DP cells. Besides Monocle, a similar developmental pathways can also be constructed by SPRING ^35^ based on the accesson matrix (Supplementary Fig. 12c). Interestingly, the pseudotime trajectory suggests three developmental pathways for this process, one of which started with stage 1 (DN) and ended in stage 2 (DP), and the other two of which started with stage 1 (DN), went through a transitional stage 3 and then bifurcated into stage 4 (CD8SP) and 5 (CD4SP). The predicted developmental trajectory could also be confirmed by the gene expression of surface markers, such as Cd4, Cd8, Runx3 and Ccr7 (Fig. 5b). To evaluate the gene ontology (GO) enrichments over the entire process, we implemented an accesson-based GO module in APEC, which highlights the significance of the association between cells and biological function (Fig. 5c). For instance, T cells selections, including β-selection, positive selection and negative selection, are initiated in the late DN stage. Consistent with this process, we observed a strong “T cell selection” GO term on the trajectory path after DN.A1 (Supplementary Fig. 12d). Since TCR signals are essential for T cell selection, we also observed the “T cell activation” GO term accompanied by “T cell selection”. Meanwhile, the signal for regulation of protein binding was found decreased at SP stages, indicating the necessity of weak TCR signal for the survival of SP T cells during negative selection.

**Figure 5.**
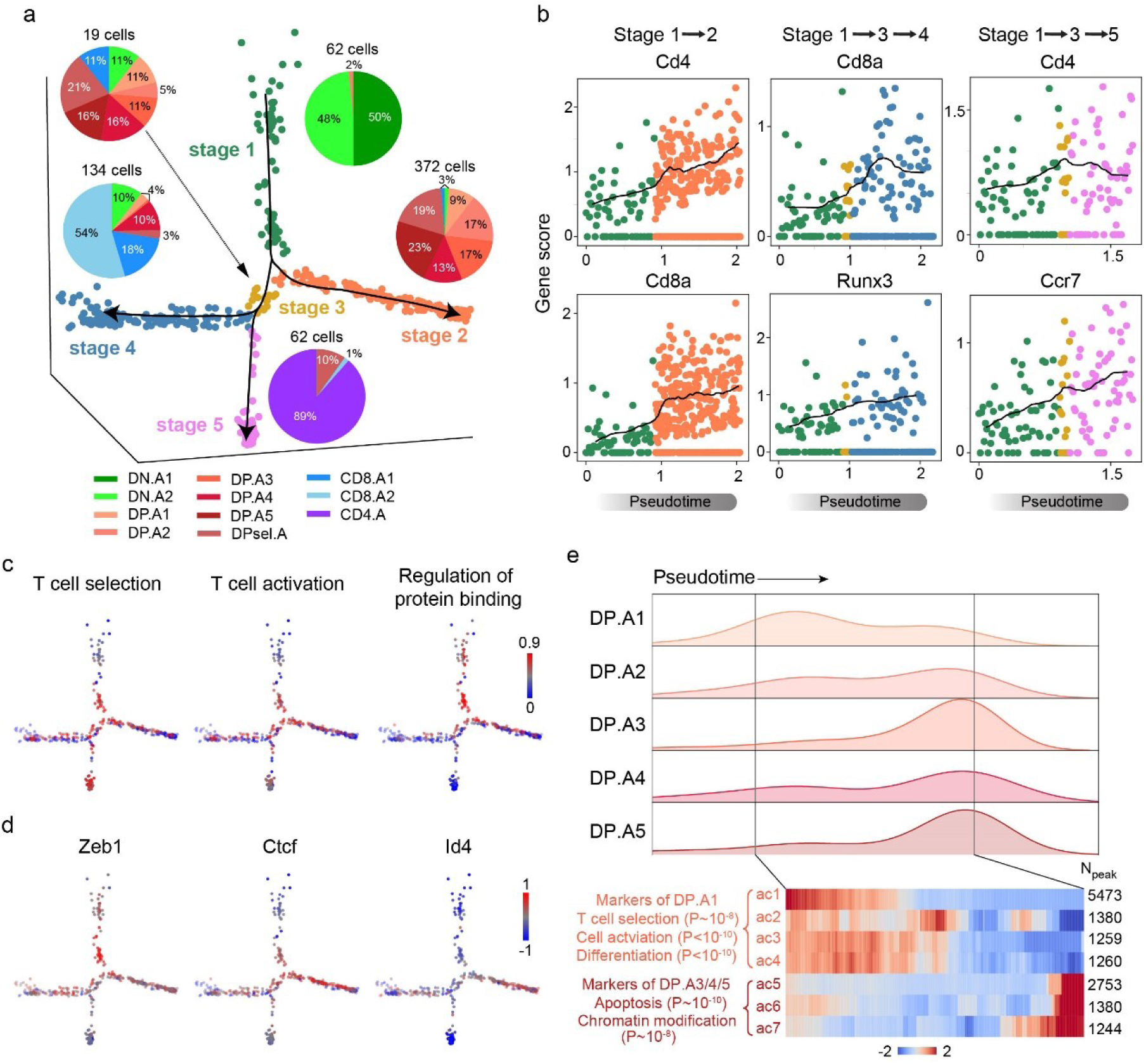
APEC depicted the developmental pathways of *Mus musculus* thymocytes by pseudotime analysis. (**a**) Pseudotime trajectory based on the accesson matrix of thymocyte ftATAC-seq data. Cell colors were defined by the developmental stages along pseudotime. Pie charts show the proportion of cell clusters at each stage. (**b**) APEC scores of important marker genes (*Cd8a*, *Cd4*, *Runx3*, and *Ccr7*) along each branch of the pseudotime trajectory. (**c**) Weighted scores of important functional GO terms along each branch of the pseudotime trajectory. (**d**) Enrichment of specific motifs searched from the differential accessons of each cell subtype. (**e**) On the stage 2 branch, the cell number distribution of clusters DP.A1∼A5 along pseudotime (upper panel) and the intensity of marker accessons of DP.A1 and DP.A3/4/5 (lower right panel) with top enriched GO terms with significance (lower left panel).

To further uncover the regulatory mechanism underlying this developmental process, APEC was implemented to identify stage-specific enriched TFs along the trajectory and pinpoint the “pseudotime” at which the regulation occurs. In addition to the well-studied TFs mentioned above (Fig. 4g), APEC also identified Zeb1 ^53^, Ctcf ^54^ and Id4 as potential stage-specific regulators (Fig. 5d). Interestingly, the Id4 motif enriched on DP cells was also reported to regulate apoptosis in other cell types ^55, 56^. Associated with the fact that the vast majority of DP thymocytes die because of a failure of positive selection ^57^, we hypothesize that stage 2 may be the path towards DP cell apoptosis. We then checked the distribution of DP cells along the stage 2 trajectory and found that most DP.A1 cells were scattered in “early” stage 2, and they were enriched with GO terms such as “T cell selection”, “cell activation” and “differentiation” (Fig. 5e, Supplementary Fig. 12e). However, most DP.A3/4/5 cells were distributed at the end of stage 2, and their principle accessons were enriched with GO terms such as “apoptosis” and “chromatin modification”. Although it is believed that more than 95% of DP thymocytes die during positive selection, only a small proportion of apoptotic cells could be detected in a snapshot of the thymus, which in our data are the cells at the end of stage 2. By comparing the number of cells near stage 3 with all the cells in stage 2, we estimated that ∼3-5% of cells would survive positive selection, which is consistent with the findings reported in previous publications ^58, 59^. Our data suggest that before entering the final apoptotic stage, DP thymocytes under selection could have already been under apoptotic stress at the chromatin level, which explains why DP cells are more susceptible to apoptosis than other thymocyte subtypes^60^.

## DISCUSSION

Here, we introduced an accesson-based algorithm for single-cell chromatin accessibility analysis. Without relying on any prior information (such as bulk sequencing data or known cell types), this approach generated more refined cell groups with reliable biological functions and properties. Integrating the new algorithm with all necessary chromatin sequencing data processing tools, APEC provides a comprehensive solution for transforming raw experimental single-cell data into final visualized results. In addition to improving the clustering of subtle cell subtypes, APEC is also capable of locating potential specific super-enhancers, searching enriched motifs, estimating gene activities, and constructing time-dependent cell developmental trajectories, and it is compatible with many existing single-cell accessibility datasets. Compared with all the other state-of-the-art single cell chromatin accessibility analysis methods, APEC clearly shows superiority in correctly predicting cell identities and precisely constructing developmental trajectories, and provides new biological insights. APEC is also very robust and stable and is scalable to clustering a large number of cells using limited computational resources. Despite these advantages, the biological implications of accessons are still obscure, especially for those that involve only a small number of peaks. Although we noticed peaks in the same accesson may belong to the same TADs, further investigations are still required to fully uncover the biology that underlies accessons.

To evaluate the performance of this approach in the context of the immune system, we adopted APEC with scATAC-seq technology to investigate the regulome dynamics of the thymic development process. Coordinated with essential cell surface markers, APEC provided a much more in-depth classification of thymocytes than the conventional DN, DP, CD4SP and CD8SP stages based on single-cell chromatin status. By reconstructing the developmental pseudotime trajectory, APEC discovered a transitional stage before thymocytes bifurcate into CD4SP and CD8SP cells and inferred that one of the stages leads to cell apoptosis. Considering that more than 95% of DP cells undergo apoptosis as a programmed cell death process, our data suggested that before DP cells enter the final apoptotic state, there would already be some intracellular changes towards apoptosis at the chromatin level. However, further studies are still needed to fully understand the regulatory mechanism of this process.

## METHODS

### ftATAC-seq on mouse thymocytes

Alexa fluor 488-labeled adaptor oligonucleotides were synthesized at Sangon Biotech as follows: Tn5ME, 5’-[phos]CTGTCTCTTATACACATCT-3’; AF488-R1, 5’-AF488-TCGTCGGCAGCGTCAGATGTGTATAAGAGACAG-3’; and AF488-R2, 5’-AF488-GTCTCGTGGGCTCGGAGATGTGTATAAGAGACAG-3’. Then, 50 μM of AF488-R1/Tn5ME and AF488-R2/Tn5ME were denatured separately in TE buffer (Qiagen) at 95 °C for 5 min and cooled down to 22 °C at 0.1 °C/s. AF488-labeled adaptors were assembled onto Robust Tn5 transposase (Robustnique) according to the user manual to form fluorescent transposomes.

Thymus tissues isolated from 6- to 8-week-old male mice were gently ground in 1 mL of RPMI-1640. Thymocytes in a single-cell suspension were counted after passing through a 40 μm nylon mesh. A total of 1 × 10^6^ thymocytes were stained with PerCP-Cy5.5-anti- CD45, PE-anti-CD8a and APC-Cy7-anti-CD4 antibodies (Biolegend) and then fixed in 1× PBS containing 1% methanal at room temperature for 5 min. After washing twice with 1× PBS, the cells were counted again. A total of 1 × 10^5^ fixed cells were resuspended in 40 μL of 1× TD buffer (5 mM Tris-HCl, pH 8.0, 5 mM MgCl_2_, and 10% DMF) containing 0.1% NP-40. Then, 10 μL of fluorescent transposomes were added and mixed gently. Fluorescent tagmentation was conducted at 55 °C for 30 min and stopped by adding 200 µL of 100 mM EDTA directly to the reaction mixture. The cells were loaded on a Sony SH800S sorter, and single cells of the CD45^+^/AF488-Tn5^hi^ population were index-sorted into each well of 384-well plates. The 384-well plates used to acquire sorted cells were loaded with 2 µL of release buffer (50 mM EDTA, 0.02% SDS) before use. After sorting, the cells in the wells were incubated for 1 min. Plates that were not processed immediately were preserved at -80 °C.

To prepare a single-cell ATAC-seq library, plates containing fluorescently tagmented cells were incubated at 55 °C for 30 min. Then, 4.2 μL of PCR round 1 buffer (1 μL of 100 μM MgCl_2_, 3 μL of 2× I-5 PCR mix [MCLAB], and 0.1 μL each of 10 μM R1 and R2 primers) were added to each well, followed by PCR: 72 °C for 10 min; 98 °C for 3 min; 10 cycles of 98 °C for 10 s, 63 °C for 30 s and 72 °C for 1 min; 72 °C for 3 min; and holding at 4 °C. Thereafter, each well received 4 µL of PCR round 2 buffer (2 μL of I-5 PCR Mix, 0.5 μL each of Ad1 and barcoded Ad2 primers, and 1 μL of ddH_2_O), and final PCR amplification was carried out: 98 °C for 3 min; 12 cycles of 98 °C for 10 s, 63 °C for 30 s and 72 °C for 1 min; 72 °C for 3 min; and holding at 4 °C. Wells containing different Ad2 barcodes were collected together and purified with a QIAquick PCR purification kit (Qiagen). Libraries were sequenced on an Illumina HiSeq X Ten system.

### SMART-seq on thymocytes

Thymocytes were stained and sorted directly into 384-well plates without fixation. SMART-seq was performed as described with some modifications ^61^. Reverse transcription and the template-switch reaction were performed at 50 °C for 1 hr with Maxima H Minus Reverse Transcriptase (Thermo Fisher); for library construction, 0.5-1 ng of cDNA was fragmented with 0.05 μL of Robust Tn5 transposome in 20 μL of TD buffer at 55 °C for 10 min, then purified with 0.8× VAHTS DNA Clean Beads (Vazyme Biotech), followed by PCR amplification with Ad1 and barcoded Ad2 primers and purification with 0.6× VAHTS DNA Clean Beads. Libraries were sequenced on an Illumina HiSeq X Ten system.

### Data source

All experimental raw data used in this paper are available online. The single-cell data for mouse thymocytes captured by the ftATAC-seq experiment can be obtained from the Genome Sequence Archive at BIG Data Center with the accession number CRA001267 and is available via http://bigd.big.ac.cn/gsa/s/yp1164Et. Other published data sets used in this study are available from NIH GEO: (1) scATAC-seq data for LSCs and leukemic blast cells from patients SU070 and SU353, LMPP cells, and monocytes from GSE74310 ^2^; (2) scATAC-seq data for HL-60 cells from GSE65360 ^8^; and (3) scATAC-seq data for hematopoietic development (HSCs, MPPs, CMPs, LMPPs, GMPs, EMPs, CLPs and pDCs) from GSE96772 ^21^. (4) APEC is also compatible with a preprocessed fragment count matrix from the snATAC-seq data for the forebrain of adult mice (p56) from GSE100033 ^10^. (5) The computational efficiency of APEC and other methods was tested using data from the single-cell atlas of mouse chromatin accessibility (sciATAC-seq) from GSE111586 ^28^. (6) The scATAC-seq (GSE65360 ^8^) and Hi-C (GSE63525 ^62^) data of GM12878 cells were used to generate the spatial correlation of peaks in the same or in different accessons.

### Preparing the fragment count matrix from the raw data

APEC adopted the general mapping, alignment, peak calling and motif searching procedures to process the scATAC-seq data from ATAC-pipe ^63^. We also implemented the python script in ATAC-pipe ^63^ to trim the adapters in the raw data (in paired-end fastq format files for each single-cell sample). APEC used BOWTIE2 to map the trimmed sequencing data to the corresponding genome index and used PICARD for the sorting, duplicate removal, and fragment length counting of the aligned data.

Unlike several previous methods that call peaks from bulk ATAC-seq data or aggregated cell populations according to cell type ^20, 21, 64^, the APEC pipeline calls peaks from the merged single-cell profiles of all cells using MACS2 to ensure that the entire analysis is unsupervised. We then ranked and filtered out the low quality peaks based on the false discovery rate (Q-value). Genomic locations of the peaks were annotated by HOMER, and motifs searched by FIMO. APEC calculates the number of fragments and the percent of reads mapped to the TSS region (±2000 BP) for each cell, and filters out high quality cells for downstream analysis. All required files for the hg19 and mm10 assembly have been integrated into the pipeline. If users want to process data from other species, they can also download corresponding reference files from the UCSC website. By combining existing tools, APEC made it possible to finish all of the above data processing steps by one command line, and generate a fragment count matrix for subsequent cell clustering and differential analysis. APEC has been made available on GitHub (https://github.com/QuKunLab/APEC).

### Accesson-based clustering algorithm

We define accesson as a set of peaks with similar accessibility patterns across all single cells, similar to the definition of gene module for RNA-seq data. After preprocessing, a filtered fragment count matrix **B** is obtained, and APEC groups peaks to construct accessons and then performs cell clustering analysis as follows:

1. *Normalization of the fragment count matrix.* Each matrix element *B*_*ij*_ represents the number of raw reads in cell *i* and peak *j*, and element *B*_*ij*_ was then normalized by the total number of reads in each cell *i*, as if there are 10,000 reads in each cell.

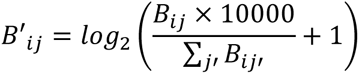
2. *Constructing accessons*. The top 40 principal components of the normalized matrix were used to construct the connectivity matrix (**C_peak_**) of peaks by the K-nearest-neighbor (KNN) method with K=10. The grouping of peaks is insensitive to the number of principal components and the number of nearest neighbors, so it is usually not necessary to change these two parameters for different data sets. Based on the matrix **C_peak_**, all peaks were grouped by agglomerative clustering with Euclidean distance and Ward linkage method, and the sum of one peak group was an accesson. For most data sets, we recommend setting the number of accessons to a value between 500 and 1500, and the default was set to 600, however the cell clustering result is not sensitive to the choice of accesson number within this range. We then built the accesson count matrix **M** by summing the fragment count of all peaks in one accesson. Thus, each column of matrix **M** is an accesson, each row is a cell, and each element represents the cumulative fragment count of each accesson in each cell. The accesson matrix was then normalized by calculating the z-score or probability of the fragment count for each row (i.e., each cell) to generate normalized matrix **M_a_** for the next step of cell clustering.
3. *Cell clustering*. From the normalized accesson matrix **M_a_**, APEC established the connectivity matrix by computing the k-neighbor graph of all cells. Since the Louvain algorithm was proven to be a reliable single-cell clustering method in Seurat ^13^ and Scanpy ^16^, we adopted it in APEC to automatically predict the number of clusters from the connectivity matrix and defined each Louvain community as a cell cluster. APEC uses the Louvain algorithm to predict cluster number to ensure that the clustering analysis is unsupervised and then performs cell clustering as default. Meanwhile, if users want to artificially define the number of cell clusters, APEC can also perform KNN clustering on the connectivity matrix.
4. *Compare the performance of APEC with that of other methods on cells with known identity*. To investigate the accuracy of the cell clusters predicted by different algorithms, we used the ARI value, which evaluates the similarity of clustering results with all known types of cells ^14^. The ARI value can be calculated as follows:

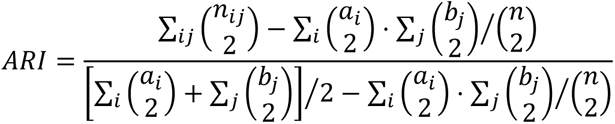

where *n*_*ij*_ is the element from the contingency matrix (i.e. the number of type *i* cells that were classified into cluster *j*), *a*_*i*_ and *b*_*j*_ are the sums of the *i* th row and *j*th column, respectively, and 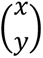 denotes a binomial coefficient. A higher ARI value indicates more accurate classification of cell types.
5. *Characteristics of accesson*. The peaks of a same accesson can be distant from each other on the genome, and sometimes even on multiple chromosomes. The average number of peaks per accesson depends on the total number of peaks in the dataset and the number of accessons set in the program (default 600). Usually the total number of peaks can vary between ∼40,000-150,000 depending on the total number of cells and the sequencing depth for each cell, thereby the average number of peaks per accesson is around ∼60-250. Beside, we chose the top 40 principle components (PCs) of the normalized matrix to construct the connectivity matrix since the first 3∼5 PCs are usually not sufficient to capture the detailed features of a single cell dataset, as described in Seurat and many other single cell analysis tools. Although default values were chosen to provide better clustering results based on analysis of multiple datasets, users can adjust these parameters as needed.

### Sampling of accesson number

To test if the APEC clustering result is sensitive to the choice of accesson numbers, we sampled 100 different accesson numbers from 500 to 1500 in steps of 10 and clustered the cells of each dataset 100 times (Fig. 1c and Supplementary Fig. 2c). APEC generated stable clustering results in terms of the average ARI on these datasets, with a wide range of different accesson numbers (Supplementary Fig. 5e). We used 600 as the default number of accessons in APEC.

### Parameter settings for other algorithms

To quantify the cell clustering performance of APEC, we compared APEC with other state-of-the-art single-cell epigenomic algorithms on the same datasets with gold standards, including cisTopic ^19^, LSI ^11, 12^, chromVAR ^17^ and Cicero ^18^. Since most of them have no cell clustering algorithm within their original codes, we applied the Louvain clustering algorithm on their transformed matrices to fairly compare their performance. We adopted the default settings of these tools for most of the comparisons in this paper, except for some parameters that were manually defined as necessary, such as the random seed in cisTopic, the number of top components in LSI, and the peak aggregation distance in Cicero. Therefore, we sampled these parameters multiple times to obtain the average ARI and ratio of correctly classified cells of the clustering results for each tool (Fig. 1c and Supplementary Fig. 2c), just as we sampled the accesson number for APEC. We set the same parameters for all the datasets as follows:

1. *cisTopic.* The scanning range of the topic number was set to [10, 40], the number of parallel CPUs was set to 5, and the random seed was sampled 100 times from 100 to 600 in steps of 5. We kept the topic matrices normalized by z-score and probability and provided the performances based on both normalization methods. We then applied the Louvain algorithm as we did in APEC to cluster cells from the normalized topic matrix generated by cisTopic.
2. *SnapATAC.* As SnapATAC uses a mapping procedure totally different with other tools, we adopt it with default parameters (binsize=5k) to build its own fragment count matrix from the merged bam file. We called the function “runJDA” with “bin.cov.zscore.lower=-2, bin.cov.zscore.upper=2, pc.num=50, norm.method =’normOVE’, max.var=5000, do.par=TRUE, ncell.chunk=1000, num.cores=1, seed.use=10, tmp.folder=tempdir()”. We then called the function “runKNN” and sampled the number of principal components from 20 to 40 (step-size=2), and the number of nearest neighbors from 10 to 20 (step-size=1). Therefore, we sampled a total of 100 times (10×10) to calculate the ARI values. Finally, we called the function “runCluster” with “louvain.lib=’leiden’, seed.use=10, resolution=1”.
3. *LSI.* We performed truncated SVD (singular value decomposition) analysis on the TF-IDF (term frequency-inverse document frequency) matrix and chose *N*_*SVD*_top components to generate the LSI matrix. *N*_*SVD*_was sampled 6 times from 6 to 11. The first component was ignored since it is always related to read depth, and the LSI scores were capped at ±1.5. Then, we used the Louvain algorithm to cluster the cells of the LSI-processed matrix.
4. *chromVAR.* The number of background iterations was set to 50, and the number of parallel CPUs was set to 1. We then used the Louvain algorithm to cluster cells based on the bias corrected deviation matrix generated by chromVAR.
5. *Cicero.* The genome window was set to 500k BPs, the normalization method was set to “log”, the number of sample regions was set to 100, the number of dimensions was set to 40, and the peak aggregation distance was sampled at 20 values from 1k to 20k BPs in steps of 1k BPs. Then, we used the Louvain algorithm to cluster cells based on the aggregated model matrix generated by Cicero.

To test the robustness of each algorithm, we randomly sampled 20%∼90% of the raw sequence reads from the dataset of AML cells and 3 cell lines (LMPP, HL60 and monocyte) and calculated the ARI accordingly. This random sampling experiment was performed 50 times for each method and average ARIs were reported (Supplementary Fig. 2d). The manually defined parameters for each method were set to: APEC, 600 accessons; cisTopic, random seed 100; SnapATAC, 20 PCs and 15 nearest neighbors; LSI, top 2-6 principle components; Cicero, 10k BPs aggregation distance.

### Gene scores and differential analysis

APEC scores a gene by the peaks around its TSS region, which is similar to the algorithm used by Preissl et al. ^10^. We calculate the average read counts of all peaks around a gene’s TSS (±20000 BP by default) as its raw score (*S*_*ij*_ for cell *i* and gene *j*), then define the gene expression by normalizing the raw score by (*S*′_*ij*_ = *S*_*ij*_ ∗ 10000⁄∑_*i*_ *S*_*ij*_), making it in a range comparable to the gene expression from scRNA-seq data. After scoring genes, APEC uses the Student’s t-test to estimate the significance of each genes between cell clusters and therefore obtained a list of differential genes filtered by P-values and fold changes. We also tested the gene scoring algorithms of other tools such as Cicero, cisTopic, and SnapATAC, and the parameters set in those algorithms were the same as described in the previous section. For scATAC-seq with very sparse signals, APEC first search for differential accessons between cell clusters, and extracted all the peaks in the differential accessons. APEC then defines genes close to these peaks (±20000 BP around TSS) as the differential genes.

### Association of cell clusters from scATAC-seq and scRNA-seq data

To determine the association between cell clusters from the epigenomics and transcriptomic sequencing, we calculated the P-values of Fisher’s exact test of the differential/non-differential genes between each pair of cell clusters from scATAC-seq and scRNA-seq data. For example, for cell cluster *a* from ftATAC-seq and cell cluster *b* from SMART-seq (Fig. 4h), if the number of consensus differential genes in both cluster *a* and *b* is G_11_, and the number of differential genes in either cluster *a* (or cluster *b*) is G_12_ (or G_21_), and the number of all the other genes is G_22_, then the 2×2 matrix **G** can be directly used for Fisher’s exact test to evaluate the P-valuie *A*_*ab*_ between cluster *a* and *b*. After constructing a matrix **A** filled with −log(*A*_*ab*_) for ftATAC-seq cluster *a* and SMART-seq cluster *b*, we calculated the z-score for each row and column of **A** to determine the correlation between cell clusters from different sequencing experiments. Same algorithm was also used to show the correspondence between the cell sub-clusters (EX1∼5 and IN1∼5) generated from snATAC-seq data and the cell subtypes identified from inDrops-seq data (Fig. 2f & g).

### Potential super-enhancers

Here, we defined a super-enhancer as a long continuous genomic area containing many accessible regions and have the same accessibility pattern in different cells. The accesson-based algorithm can group most peaks in one super-enhancer to one accesson since they always present the same accessibility pattern between cells. APEC identified super-enhancers by counting the number of peaks in a 1 million BP genomic area that belong to a same accesson. It also requires that the percentage of the putative peaks in one super-enhancer is one of the highest among all genomic areas (P-value<0.01). The pipeline can also aggregate bam files by cell types/clusters and convert them to BigWig format for users to upload to the UCSC genome browser for visualization. ROSE ^27^ was used to confirm the super enhancers called from APEC, with command line “python ROSE_main.py -g hg19 -i top_filtered_peaks.gff -r P1-LSC.bam -s 12500 -t 2500 -o P1-LSC-SuperEnhancer” applied to the merged bam files of all the P1-LSC cells.

### Spatial correlation of peaks in the same accesson

To test whether peaks in the same accesson are closer in space, we integrated the Hi-C ^62^ data (GSE63525) and scATAC-seq ^8^ data (GSE65360) on GM12878 cells. The spatial correlation of different windows, both intra- and inter-chromosomal, can be directly extracted by Juicer ^65^. The Pearson’s correlation matrix of the intra-chromosomal or inter-chromosomal windows can be calculated from the corresponding observed/expected matrix. We constructed 600 accessons by grouping peaks in the GM12878 scATAC-seq data in APEC, and removed the accessons that contained more than 1000 peaks or less than 5 peaks. The width of the window was set to 500k BPs, and peaks were then assigned to each window. Next, we collected Hi-C correlations between windows that contained peaks in the same accesson, termed “Accesson” correlations. For comparison, we also shuffled all peaks in different accessons to make fake accessons and re-collected the Hi-C correlations between windows that contained peaks in each fake accesson, termed “Shuffled” correlations. Meanwhile, we also collected Hi-C correlations between windows that contained no peaks, termed “Non-accesson” correlations. We made boxplots for these three types of correlations for intra-chromosomal and inter-chromosomal peaks and found that co-accessible peaks are spatially closer to each other than random peaks.

### Pseudotime trajectory constructed by APEC

As a tool to simulate the time-dependent variation of gene expression and the cell development pathway, Monocle has been widely used for the analysis of single-cell RNA-seq experiments ^32, 66^. APEC reduced the dimension of the accesson count matrix **M** by PCA, and then performed pseudotime analysis using the Monocle program. For complex datasets, it is necessary to limit the number of principal components, since too many features will cause too many branches on the pseudotime trajectory, and makes it difficult for a user to identify the biological significance of each branch. For the hematopoietic single cell data and thymocyte data, we used the top 5 principal components of the accesson matrix to construct the developmental and differentiation trajectories.

### Pseudotime trajectory constructed by other algorithms

To check whether other algorithms can provide solutions to construct cell developmental pathways, we combined their transformed count matrix with Monocle to build the pseudotime trajectory from scATAC-seq data. A similar preprocessing method was applied to ensure the fairness of the comparison:

1. Raw fragment count matrix. We normalized the raw count matrix **B** exactly as in the first step of APEC, i.e., 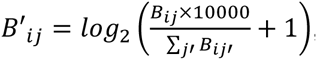, and performed PCA analysis on the normalized matrix B’. Only the top 5 PCs were subjected to Monocle to construct the trajectory.
2. cisTopic. The topic matrix generated by cisTopic was normalized by making the sum of each row the same (i.e., the probability). Then, we performed PCA analysis on the normalized topic matrix and subjected the top 5 PCs to Monocle to build the trajectory.
3. SnapATAC. We run PCA analysis on the normalized fragment count matrix generated by SnapATAC, and subjected the top 5 PCs to Monocle to build the trajectory.
4. LSI. We chose the 2nd∼6th principle components of the SVD transformation of the LSI matrix and subjected them to Monocle to construct the trajectory. As the dimensions had been reduced by the LSI method, we skipped the PCA analysis.
5. ChromVAR. After the PCA analysis of the bias corrected deviation matrix generated by chromVAR, the top 5 PCs were combined with Monocle to construct the trajectory.
6. Cicero. We performed PCA analysis on the aggregated matrix generated by Cicero and used the top 5 PCs in Monocle to build the trajectory.

In addition, to confirm the reliability of the APEC + Monocle prediction of the developmental pathway, we applied another pseudotime trajectory constructing method, SPRING ^35^, to the accesson count matrix **M** from APEC to reconstruct the pathways for the hematopoietic differentiation dataset and thymocyte developmental dataset. We performed PCA analysis of the accesson matrix **M** and subjected the top 5 PCs to SPRING to generate the trajectories. The number of edges per node in SPRING was set to 5.

### Parameter settings for each dataset

In the quality control (QC) step, cells are filtered by two constraints: the percentage of the fragments in peaks (*P*_*f*_) and the total number of valid fragments (*N*_*f*_). However, there is no fixed cutoff for these two parameters since the quality of different cell types and/or experiment batches are completely different. The total number of peaks is usually limited to approximately 50000 to reduce computer time, but we recommend using all peaks if the users want to obtain better cell clusters. (1) For the data set from hematopoietic cells, the -log(Q-value) threshold of high-quality peaks was set to 35 to retain 54212 peaks, and the cutoff values of *P*_*f*_ and *N*_*f*_ were 0.1 and 1000, respectively. (2) For the scATAC-seq data on the two types of cells from 2 AML patients (P1-LSC, P1-Blast, P2-LSC, P2-Blast), the threshold of -log(Q-value) was set to 5 to retain 38683 high-quality peaks for subsequent processing. When LMPPs, HL60 and monocytes were added to this dataset with the AML cells, the threshold of -log(Q-value) was set to 8 to retain 42139 peaks. In the QC step, we set the *P*_*f*_ cutoff to 0.05 and the *N*_*f*_ cutoff to 800. (3) For the snATAC-seq data from the adult mouse forebrain, all peaks and the raw count matrix obtained from the original data source were adopted in the analysis. (4) For the ftATAC-seq data from thymocytes, all 130685 peaks called by MACS2 were reserved for the fragment count matrix (Q-value<0.05), and we retained cells with *P*_*f*_ >0.2 and *N*_*f*_ >2000.

### SMART-seq data analysis with Seurat

For the analysis of SMART-seq data from mouse thymocytes, we employed STAR (version 2.5.2a) with the ratio of mismatches to mapped length (outFilterMismatchNoverLmax) less than or equal to 0.05, translated output alignments into transcript coordinates (i.e., quantMode TranscriptomeSAM) for mapping ^67^ and used RSEM ^68^ to calculate the TPM of genes. For QC, we excluded cells in which fewer than 2000 genes were detected and genes that were expressed in only 3 or fewer cells. Seurat filtered cells with several specific parameters to limit the number of genes detected in each cell to 2000∼6000 and the proportion of mitochondrial genes in each cell was set to less than 0.4 (i.e., low.thresholds=c(2000, Inf), high.thresholds=c(6000, 0.4)). Additionally, the top 12 principal components were used for dimension reduction with a resolution of 3.2 (dims.use =1:12, resolution=3.2), followed by cell clustering and differential expressed gene analysis ^69^.

### GO term analysis of cells along pseudotime trajectory

We defined the functional characteristics of each accesson by the GO terms and motifs enriched on its peaks. The GO terms of an accesson were obtained by submitting all of its peaks to the GREAT website ^70^. The negative logarithm of the P-value of each GO term in each accesson was filled into a (GO terms) × (accessons) matrix **L**. The significance of each GO term on each cell was evaluated by the product of the matrix **L** and the accesson count matrix **M**, i.e.

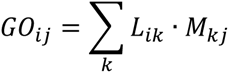

where *i* is the *i*th GO term, *j* is the *j*th cell, and *k* is the *k*th accesson. Then we calculated the z-score for each row of this product matrix, and plotted the z-score as the GO-term score on the trajectory diagram.

### Motif enrichment of cells along pseudotime trajectory

To assess the motif enrichment of the accessons, we used the Centrimo tool of the MEME suite ^71^ to search for the enriched motifs for the peaks of each accesson and applied the same algorithm as to the GO term score to obtain the motif score. The negative logarithm of the E-value (product of adjusted P-value and motif number) ^71^ of each motif in each accesson was used to construct a (motifs) ×(accessons) matrix **F**. The enrichment of each motif on each cell was evaluated by the product of the matrix **F** and the accesson count matrix **M**, i.e.,

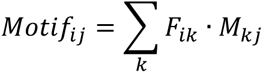

where *i* is the *i*th motif, *j* is the *j*th cell, and *k* is the *k*th accesson. Then, we calculated the z-score for each row of this product matrix and plotted the z-score as the motif score on the trajectory diagram.

## DATA AND CODE AVAILABILITY

Mouse thymocytes ftATAC-seq data can be obtained from the Genome Sequence Archive at BIG Data Center with the accession number CRA001267 and is available via http://bigd.big.ac.cn/gsa/s/yp1164Et. Other published data sets used in this study are available from NIH GEO with accession numbers GSE74310 ^2^, GSE65360 ^8^, GSE96772 ^21^, GSE100033 ^10^, GSE111586 ^28^, and GSE63525 ^62^. APEC pipeline can be downloaded from the GitHub website (https://github.com/QuKunLab/APEC).

## ACKNOWLEDGMENTS

This work was supported by the National Key R&D Program of China (2017YFA0102900 to K.Q.) and by National Natural Science Foundation of China grants (81788101, 91640113, 91940306, 31771428 to K.Q.). It was also supported by the Fundamental Research Funds for the Central Universities (to K.Q.) and the Anhui Provincial Natural Science Foundation grant BJ2070000097 (to B.L.) and 1908085QH326 (to Y.L.). We thank the Howard Chang lab at Stanford University for helpful discussion. We thank the USTC supercomputing center and the School of Life Science Bioinformatics Center for providing supercomputing resources for this project.

## AUTHOR CONTRIBUTIONS

KQ, BL and YL conceived the project, BL developed the APEC software and performed all data analysis with helps from KL, QY, PC, JF, WZ, PD and CJ. YL developed ftATAC-seq technique and performed all scATAC-seq and scRNA-seq experiments with helps from LZ. KL analyzed scRNA-seq data. BL, YL and KQ wrote the manuscript with inputs from all other authors.

## Supplementary Figures

**Supplementary Fig. 1.**
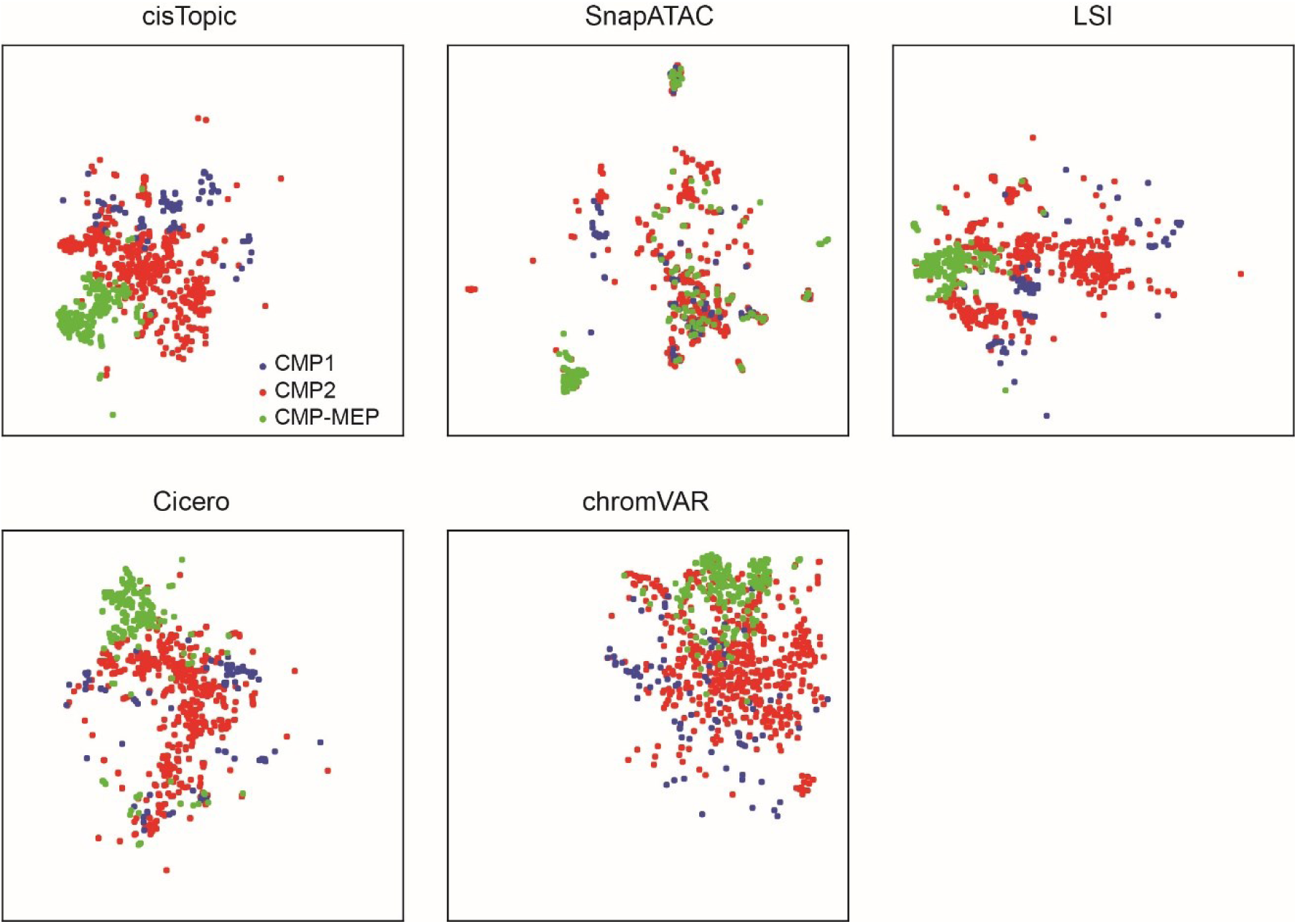
The 3 subtypes of CMP cells in the tSNE maps generated by other methods.

**Supplementary Fig. 2.**
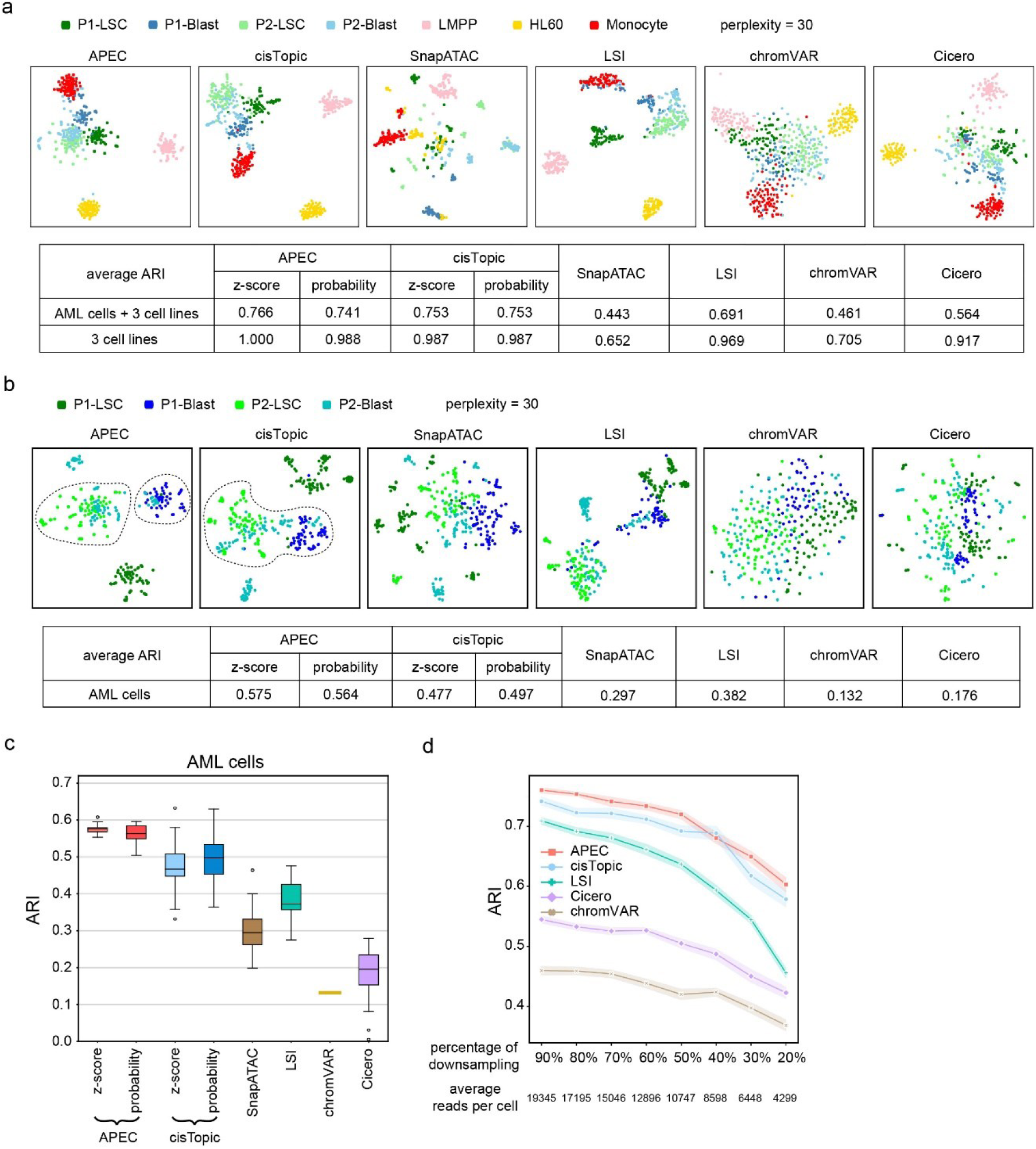
Clustering performance of the dimension-transformed matrices generated by different algorithms. (**a**) The tSNE diagrams of the cells from AML patients and three distinct cell lines (LMPP, monocyte and HL60). Different algorithms provided different dimension-transformed matrices for tSNE analysis, i.e., APEC: accesson matrix; cisTopic: topic matrix; LSI: LSI matrix; chromVAR: bias corrected deviation matrix; Cicero: aggregated model matrix. The table below the diagrams contains the average ARI of the cell clustering results for each algorithm. (**b**) The tSNE diagrams and ARI table for the leukemic stem cells (LSCs) and blast cells from 2 different AML patients only, as in (a). (**c**) Box-plots showing the ARI values for the clustering of the blast and LSC cells from two AML patients. We sampled different tunable parameters for different algorithms. APEC: the accesson number; cisTopic: the random seed; SnapATAC: the number of principal components and the number of nearest neighbors; LSI: the number of top SVD components; Cicero: the peak aggregation distance; chromVAR: no sampling. Z-score and probability denote different methods of normalizing the dimension-transformed matrices. Center line, median; box limits, upper and lower quartiles; whiskers, 1.5x interquartile range; points, outliers. (**d**) The average ARI values calculated by down-sampling 50 times from the raw data of the AML cells and three cell lines for each method. The X-axis represents the percentage of down-sampled sequencing reads. Shaded error band: 95% confidence interval.

**Supplementary Fig. 3.**
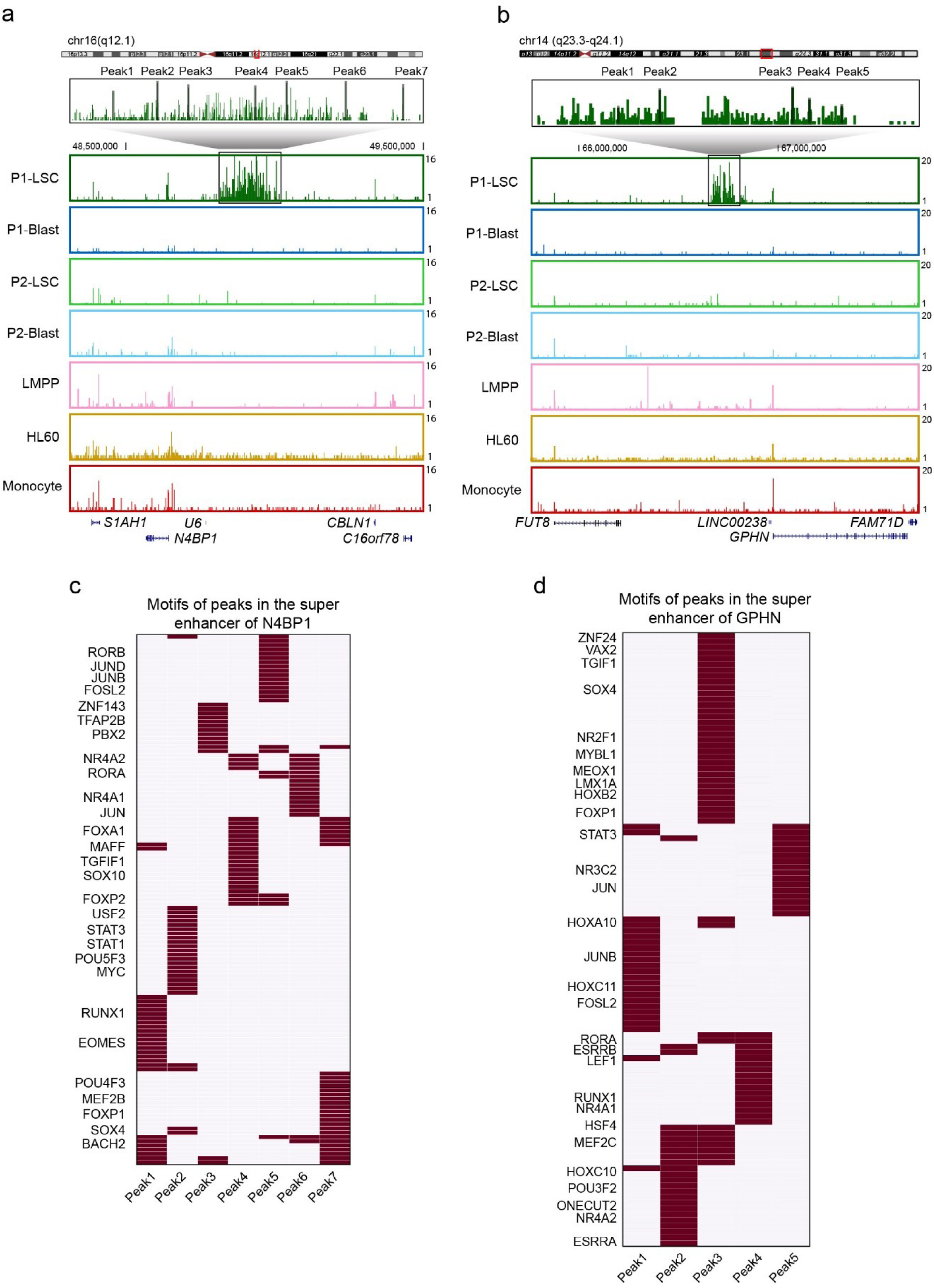
Super-enhancers predicted by APEC for the scATAC-seq data of cells from AML patients. (**a, b**) The genome browser track shows the aggregated scATAC-seq signal of the super-enhancer of P1-LSC cells upstream of *N4BP1* (a) and *GPHN* (b). (**c, d**) The motifs associated with peaks in the super-enhancer upstream of *N4BP1* (c) and *GPHN* (d).

**Supplementary Fig. 4.**
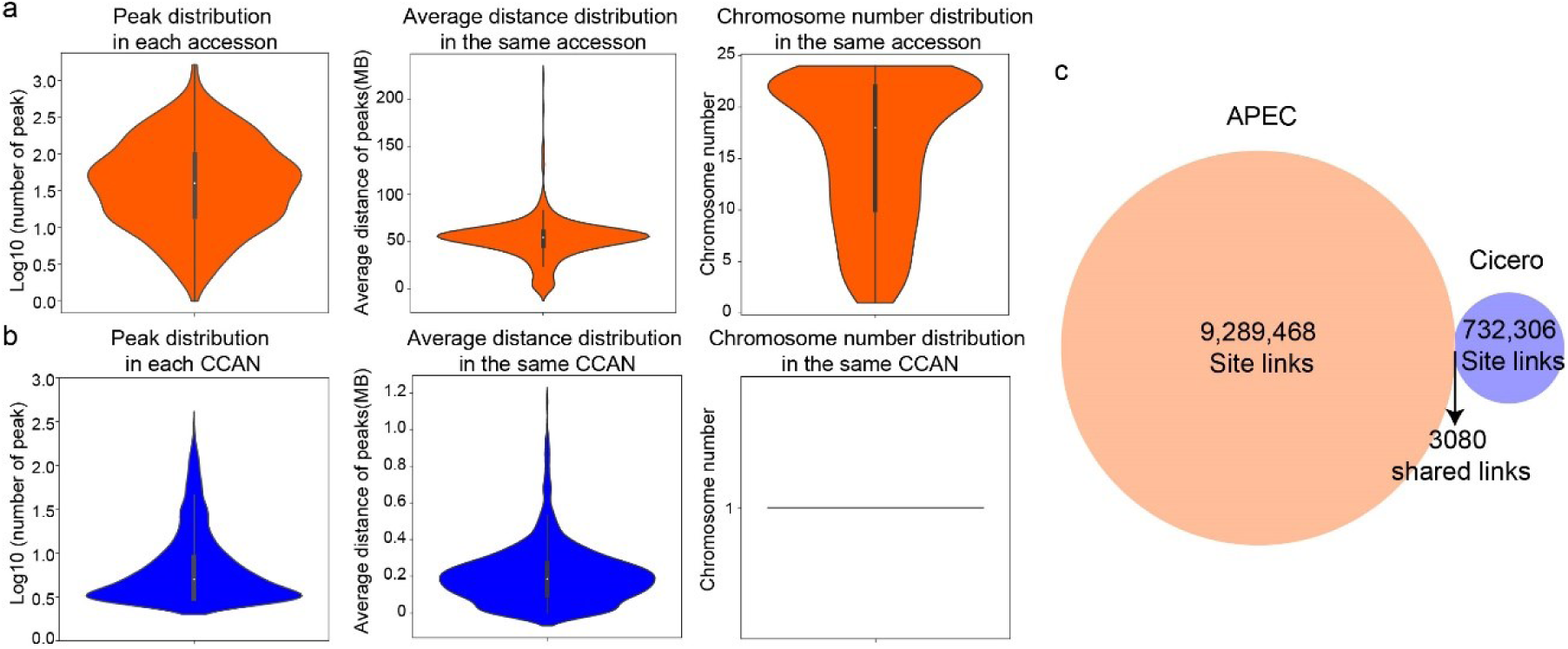
Comparison of the peak grouping algorithms used by APEC and Cicero on the hematopoietic dataset. (**a**) The characteristics of accessons in APEC. Left panel: distribution of peaks in each accesson; middle panel: genomic distances of peaks belong to the same accesson; right panel: number of chromosomes with peaks belong to the same accesson. (**b**) The characteristics of CCAN (defined by Cicero), as in (a). (**c**) Site links discovered by APEC and Cicero.

**Supplementary Fig. 5.**
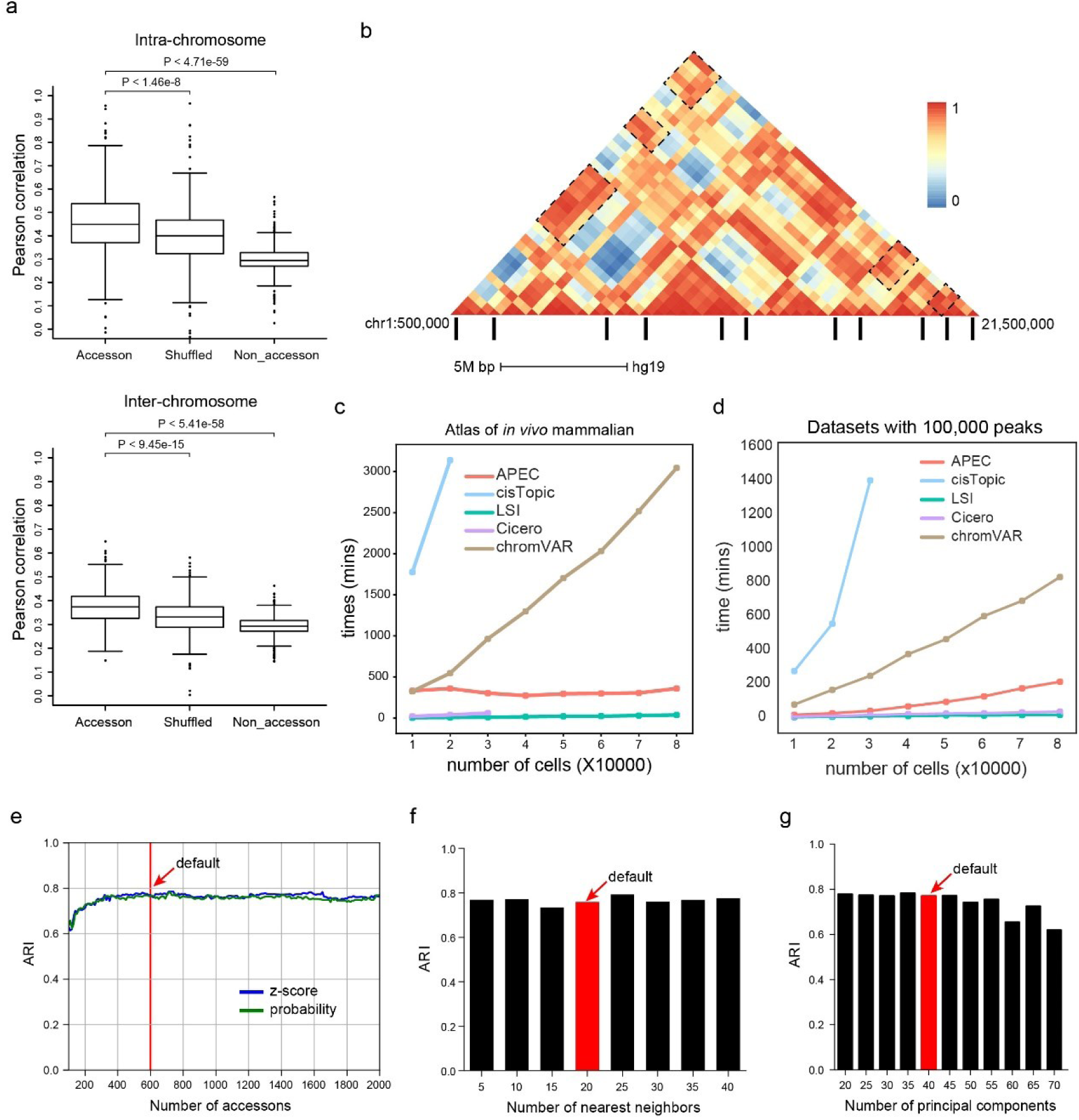
Biological insights of the accesson and stability and scalability analysis of APEC. (**a**) Box plot showing the average spatial distance between peaks in the same accesson (from scATAC-seq) versus randomly shuffled peaks versus non-accessible genomic regions. Spatial distance was estimated from chromosome conformation capture (Hi-C) technology. Both Hi-C and scATAC-seq data were generated from the same cell line GM12878. Upper panel: intra-chromosomal correlation of windows in the Hi-C data; Lower panel: inter-chromosomal correlation of windows in the Hi-C data. Accesson: The correlation between two windows that contain peaks in the same accesson; Shuffled: The correlation calculated by randomly shuffling peaks in each accesson; Non-accesson: The correlation between two windows that contain no peaks. Center line, median; box limits, upper and lower quartiles; whiskers, 1.5x interquartile range; points, outliers. (**b**) The Hi-C profile of windows between chr1:500,000-21,500,000. The black bars below the Hi-C track denote peaks in the same accesson from APEC. Dotted boxes indicate examples of peaks in the same accesson that are distant in genomic position but close in space. (**c, d**) The computing time required for different algorithms to cluster cell numbers from 10,000 to 80,000 with all peaks (c) and 100,000 peaks (d). The data were sampled from the single-cell atlas of *in vivo* mammalian chromatin accessibility. CisTopic was performed using 8 CPU threads and all the other tools with 1 CPU thread. (**e-g**) The ARI values of the clustering results that used different numbers of accessons (e), nearest neighbors (f), and principle components (g). The dataset includes the cells from two AML patients and three cell lines. Default values are noted in red.

**Supplementary Fig. 6.**
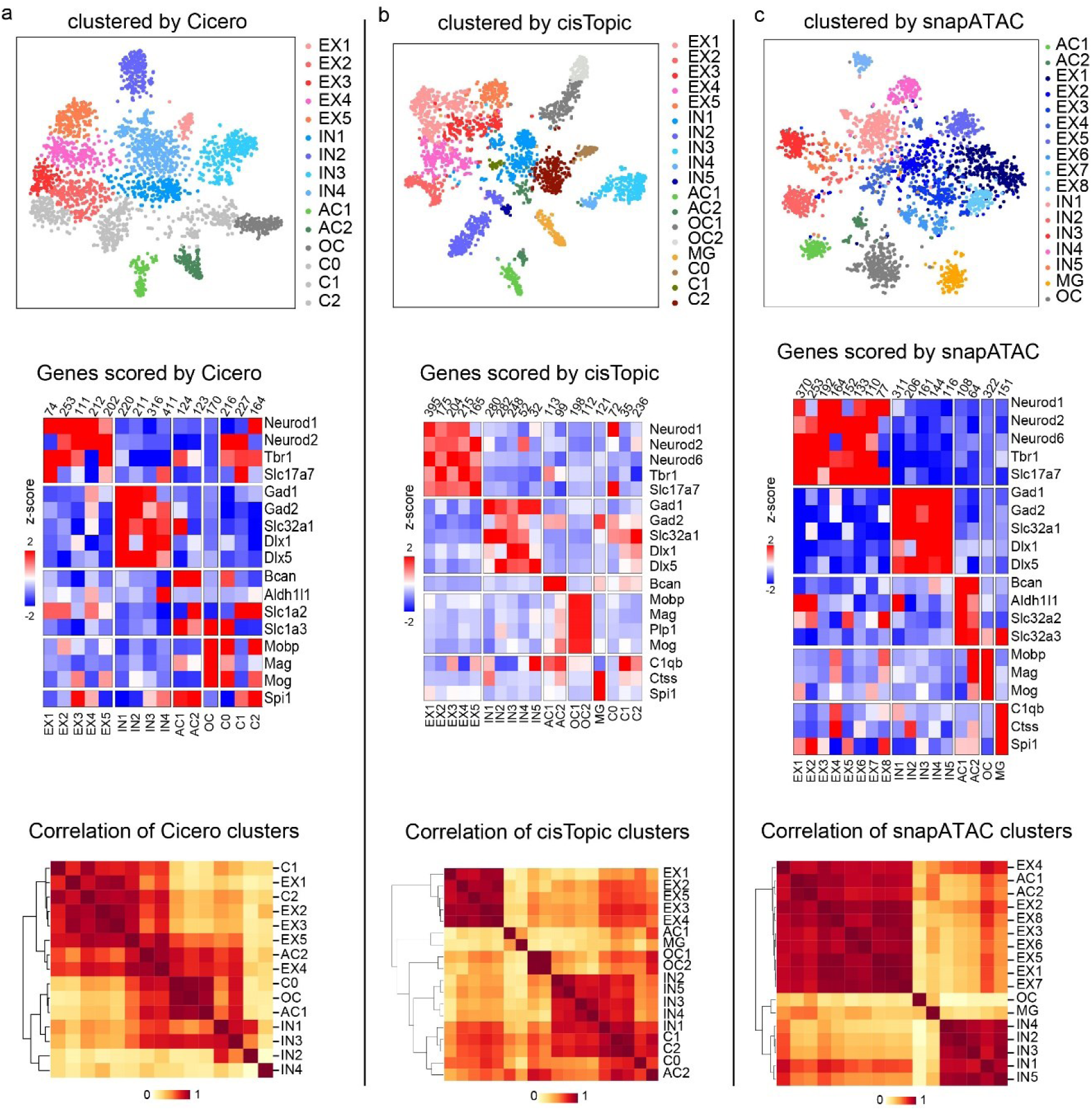
(**a**) The clustering and cell-type classification of the mouse forebrain dataset by Cicero. Upper panel: cell clusters obtained by Cicero, illustrated in the tSNE diagram. Middle panel: the z-scores of the average gene scores of cell clusters, obtained by Cicero. Lower panel: the hierarchical clustering of the Pearson correlations between cell clusters identified by Cicero. (**b, c**) The clustering and cell-type classification of the same dataset by cisTopic and SnapATAC respectively, as in (a).

**Supplementary Fig. 7.**
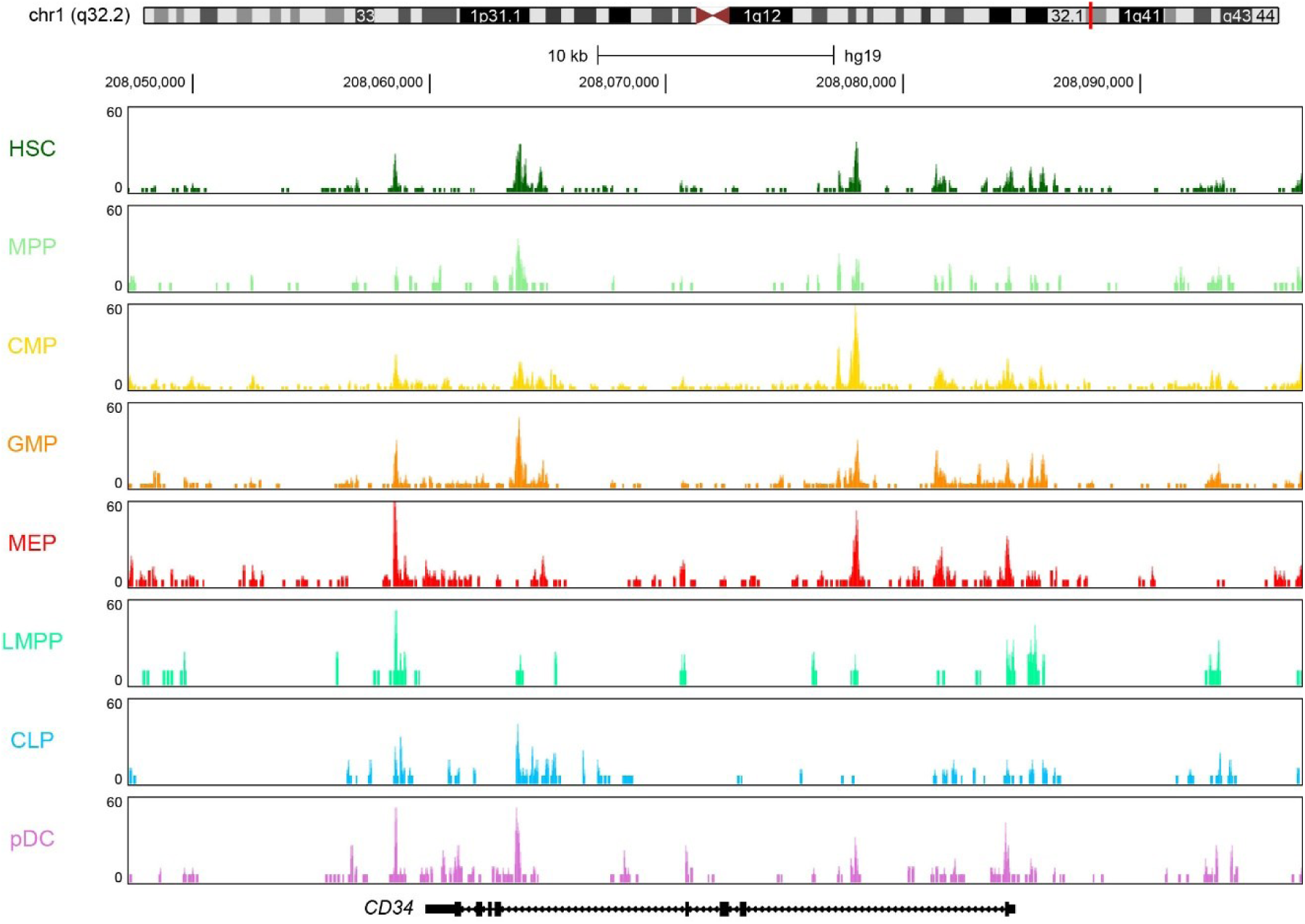
UCSC genome browser track diagram of the normalized fragment count around gene *CD34* for each hematopoietic cell type.

**Supplementary Fig. 8.**
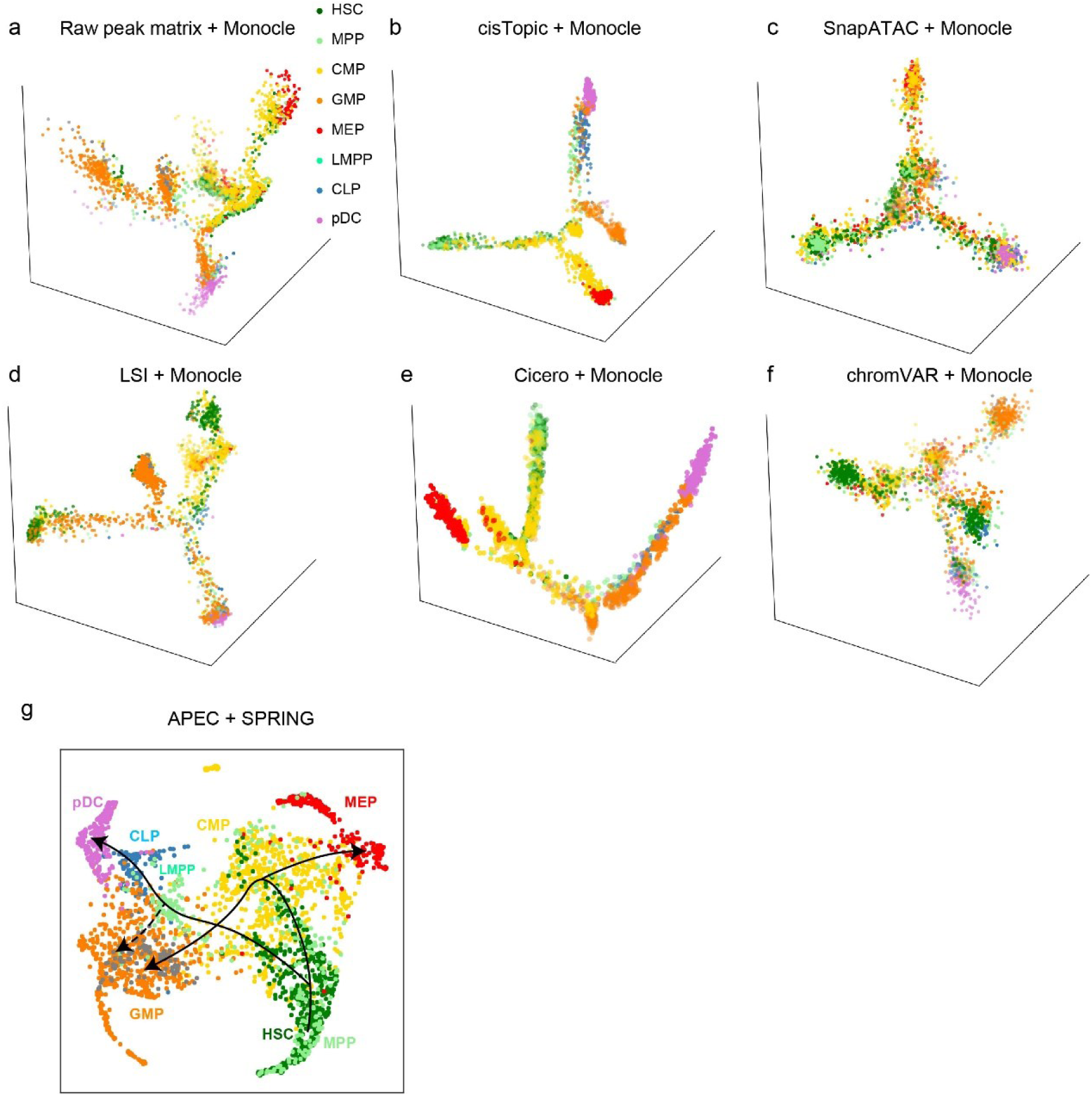
Cell differentiation trajectories of the human hematopoietic dataset constructed by different algorithms. (**a-f**) The pseudotime trajectories constructed by the combination of Monocle and the raw peak count matrix, the topic matrix from cisTopic, the normalized count matrix from SnapATAC, the LSI matrix, the aggregated model matrix from Cicero, and the bias corrected deviation matrix from chromVAR, respectively. (**g**) The pseudotime trajectory constructed by the combination of SPRING and the accesson matrix from APEC.

**Supplementary Fig. 9.**
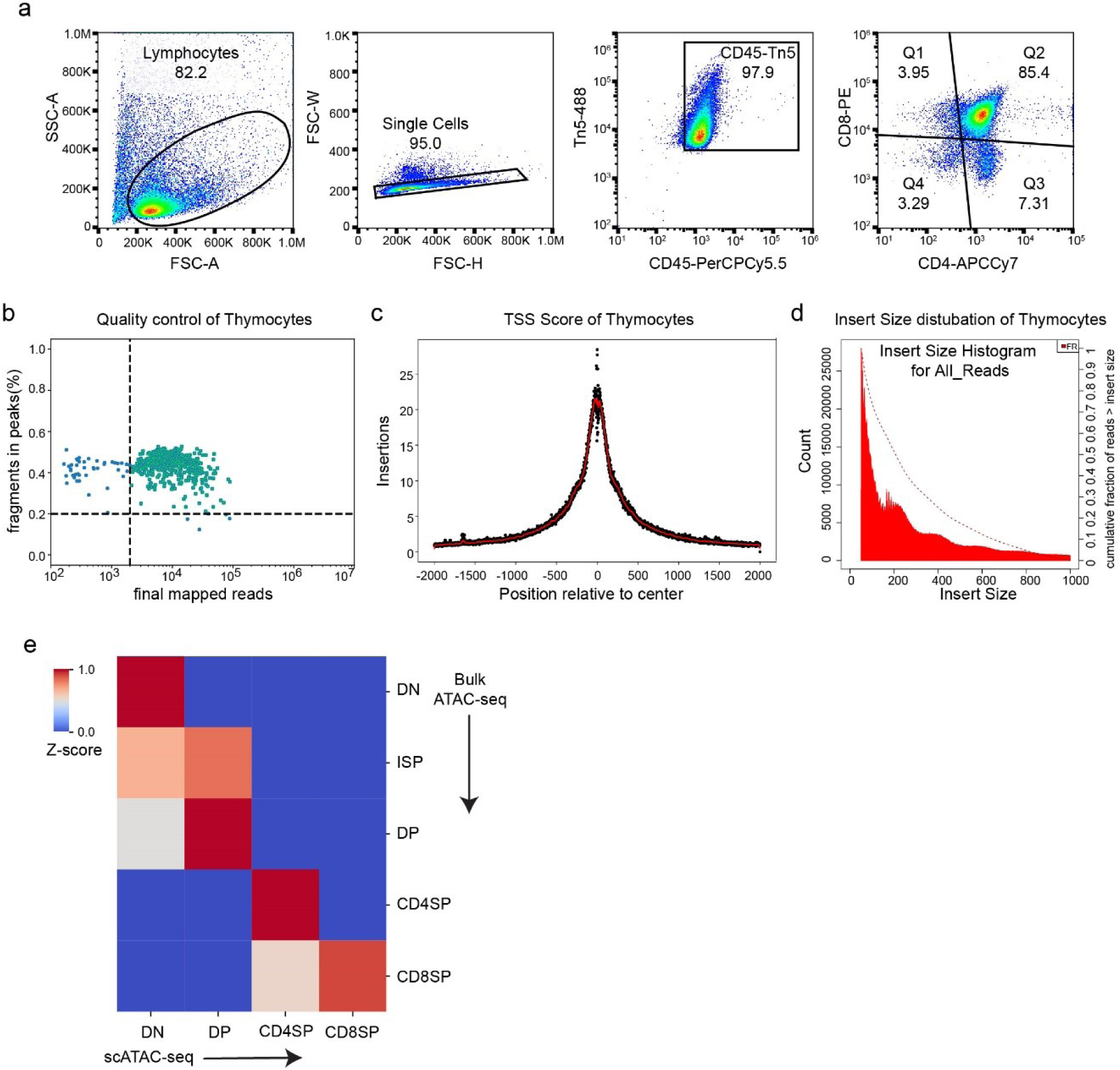
(**a**) Gating strategy of the mouse thymocytes in ftATAC-seq. (**b-d**) Quality control diagrams for the mouse thymocyte data, including reads numbers and percentage of fragments in peaks for each cell (b), average count of scATAC-seq insertions around TSS regions (c), and statistical distribution of fragment lengths (d). (**e**) The z-score of correlation between the cell types from ftATAC-seq and bulk ATAC-seq data.

**Supplementary Fig. 10.**
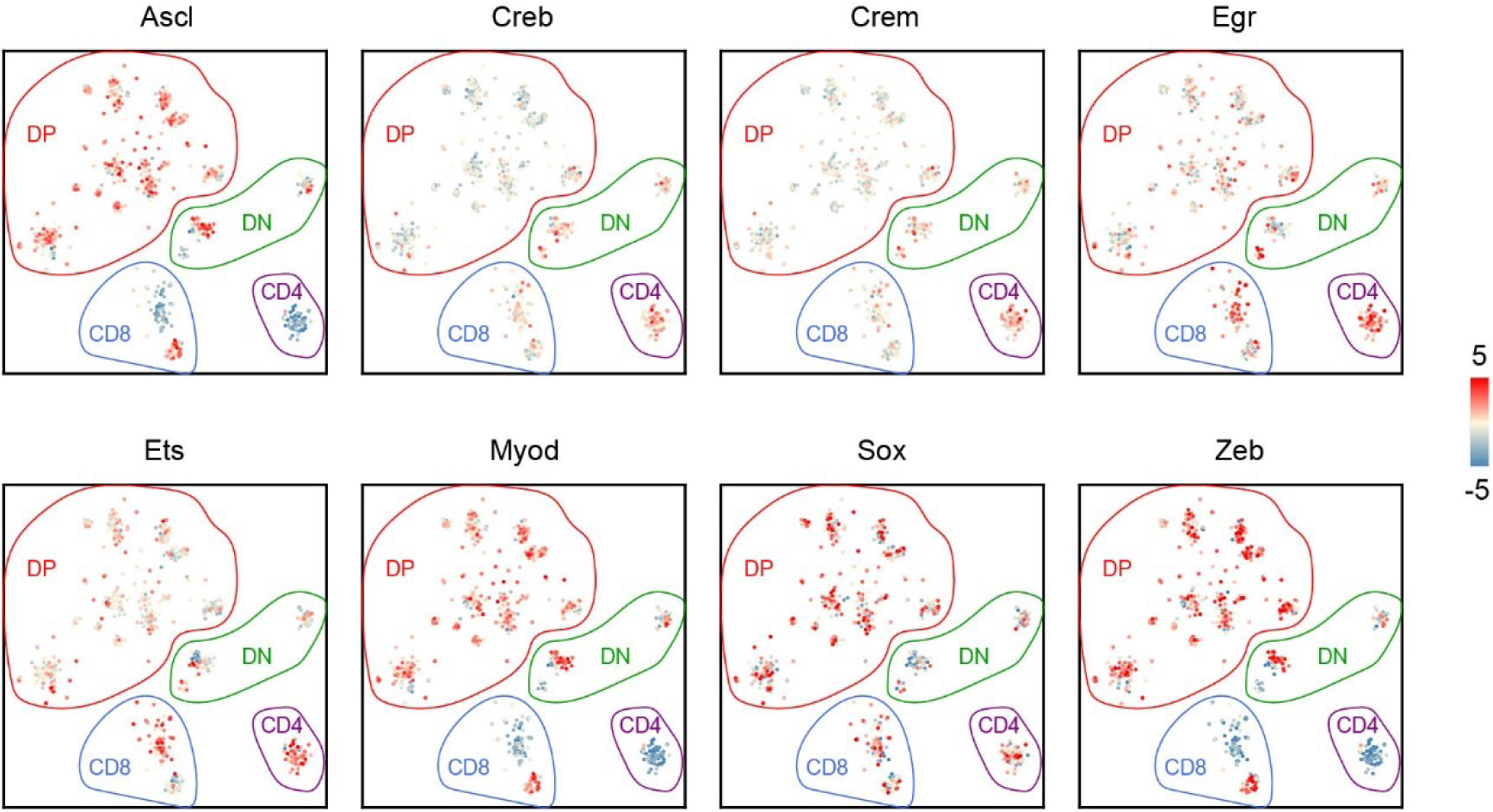
Selected significant motifs enriched in different thymocyte subtypes obtained by the APEC algorithm.

**Supplementary Fig. 11.**
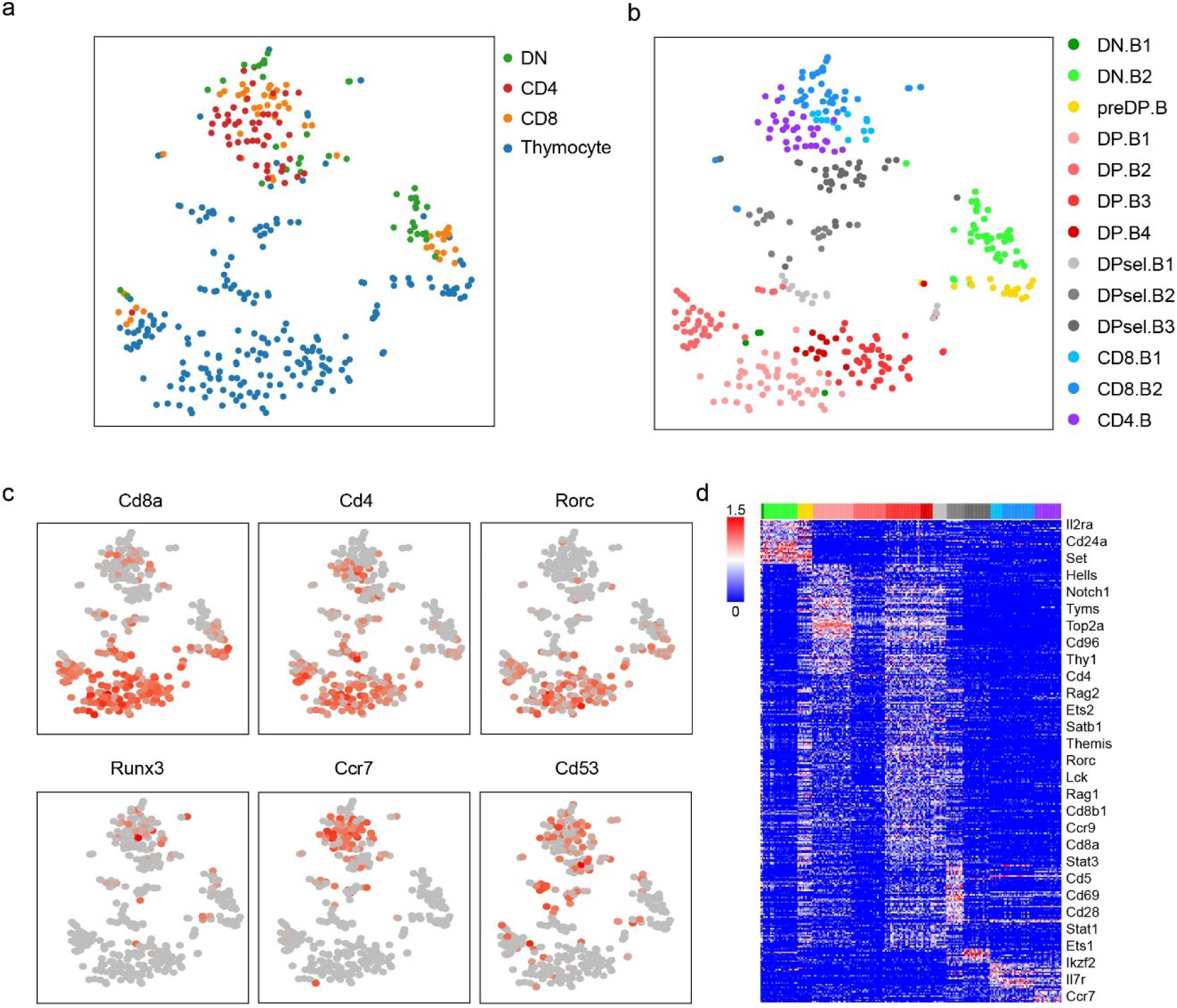
Single-cell transcriptome analysis of *Mus musculus* thymocytes from SMART-seq. (**a**) tSNE diagram of the single-cell expression matrix of *Mus musculus* thymocytes, labeled by the FACS index of each cell. (**b**) Louvain clustering of the same single-cell dataset obtained by Seurat. The cell types of these clusters were classified by the expression of corresponding marker genes. (**c**) Important marker genes were differentially expressed in different cell clusters. (**d**) Heatmap of the expressions of all genes significantly differentially expressed between cell clusters. The top color bar used the same scheme described in (b) to render cells of different clusters.

**Supplementary Fig. 12.**
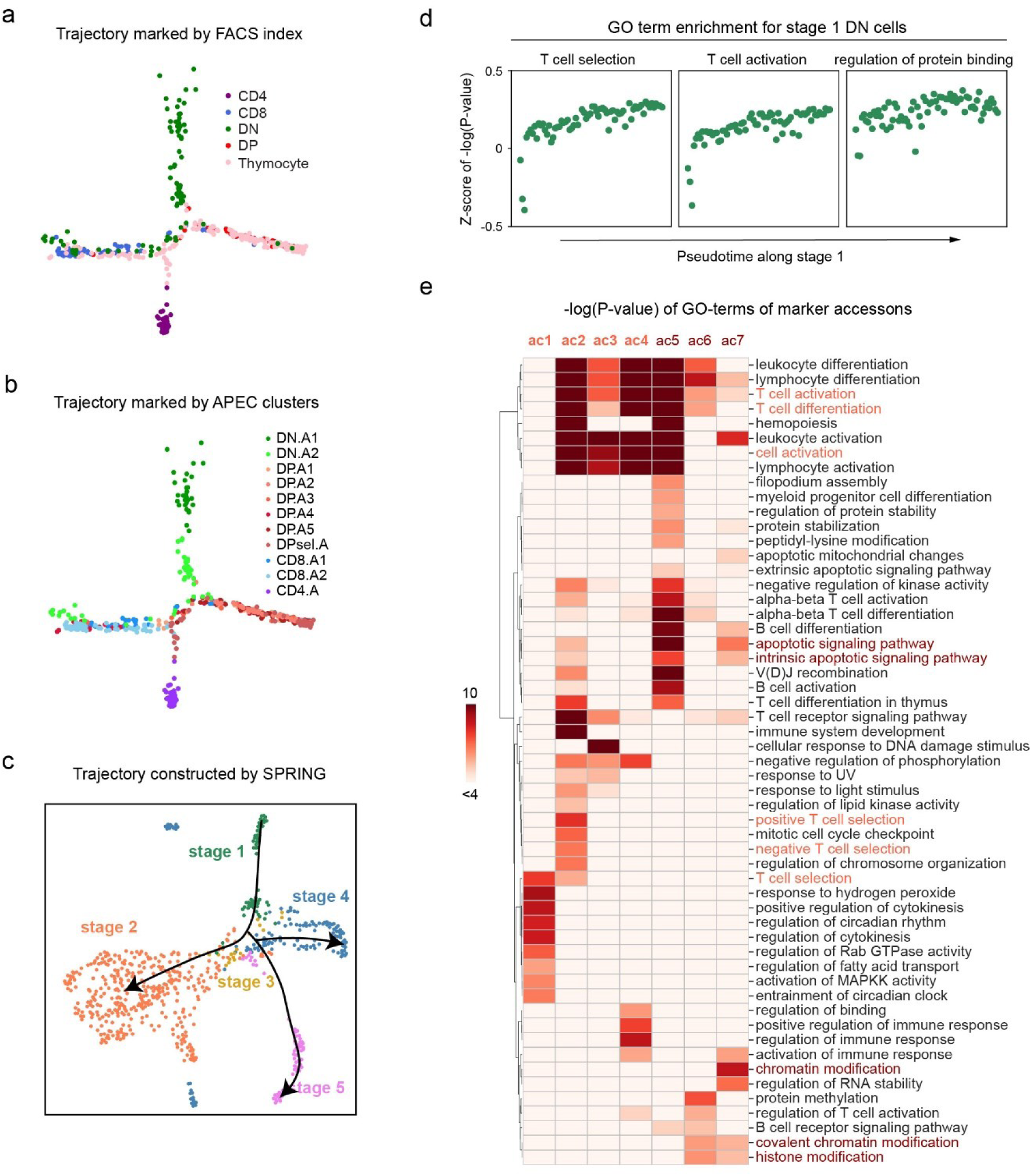
Developmental characteristics of single-cell samples captured by APEC. (**a, b**) Pseudotime trajectory of scATAC-seq data from *Mus musculus* thymocytes labeled with the FACS index and APEC cluster index. (**c**) Pseudotime trajectory constructed by applying SPRING to the accesson matrix. The colors of cells denote their stages in the APEC trajectory results. (**d**) Z-scores of the -log(P-value) of the GO terms along the pseudotime trajectory of stage 1 cells. (**e**) Logarithm of the P-value of GO terms searched from peaks in accessons ac1∼ac7, which are the marker accessons of cluster DP. A1 and DP. A3/4/5 of thymocytes.

